# LAP-TLR2 Axis Drives Inflammatory Pathology during *Listeria monocytogenes* Gastroenteritis

**DOI:** 10.64898/2026.11.05.764615

**Authors:** Dongqi Liu, Shivendra Tenguria, Nicole Irizarry-Tardi, Joonyeop Yeo Franklin, Sungtae Park, Nicholas L.F. Gallina, Luping Xu, Vignesh Nathan, Xilin Li, Yang Fu, Rishi Drolia, Xingjian Bai, Zihan Gao, Lavanya Reddivari, Kurt D. Ristroph, Abigail Cox, Matthew R. Olson, Arun K. Bhunia

## Abstract

*Listeria monocytogenes* (Lm) causes acute gastroenteritis that precedes systemic invasive listeriosis, yet the molecular mechanisms coordinating bacterial virulence with host inflammatory responses during the enteric phase remain poorly understood. We show that Lm-associated gastroenteritis is dose-dependent, and a high infectious dose (∼1×10^10^ CFU/mouse), reflecting epidemic cases, recapitulates hallmark features of gastroenteritis in a mouse model, including early onset, epithelial erosion, and profound inflammatory cell infiltration. We identify *Listeria* adhesion protein (LAP), rather than classical virulence factors such as internalin A and listeriolysin O, as the primary mediator of enteric pathology, driving epithelial cell death and neutrophil recruitment. Transcriptomic (mRNA-seq) profiling implicates LAP-driven engagement of the TLR2-MyD88 pathway. Using the HEK-Dual™ hTLR2 (NF-κB) reporter cell line and *TLR2* knockout mice, we show that LAP engages TLR2 to activate TNFα–NF-κB signaling and compromise the gut epithelial barrier. Disease severity correlates with myeloid hyperactivation and intestinal ion channel dysregulation, as *CD11b⁻^/^⁻* mice exhibit attenuated diarrhea despite elevated bacterial tissue loads. Finally, studies in *MLCK* knock-out mice confirm that this myeloid-driven pathology requires MLCK-dependent tight junction opening for paracellular neutrophil migration, which works in concert with LAP–TLR2 signaling during Lm-induced gastroenteritis.

## Introduction

Gastroenteritis is characterized by epithelial barrier dysfunction, dysregulation of ion transport, and infiltration of inflammatory cells, which can culminate in erosion or loss of the mucosal epithelial layer ^1^. Diarrhea is one of the hallmarks of acute infectious gastroenteritis, manifesting as watery or bloody, depending on the mechanism of the pathophysiology. Although gastroenteritis is commonly associated with many enteric pathogens, *Listeria monocytogenes* (Lm) is an underappreciated cause of acute febrile gastroenteritis ^2–6^. While typically self-limiting in healthy individuals, Lm-induced gastroenteritis can be life-threatening in immunocompromised populations, including children, the elderly, and pregnant women ^7,8^. In addition to immunosuppressive conditions, gastric acid-suppressing drugs, such as H_2_ blockers, are significant risk factors for febrile gastroenteritis ^9^.

Lm-induced gastroenteritis may be underrecognized in clinical settings because Lm is more commonly associated with neurological complications than with gastrointestinal disease. Recent epidemiological trends indicate that in high-risk populations, gastroenteritis often precedes fatal invasive neurolisteriosis or maternal-fetal listeriosis ^7,8,10–14^. Historical outbreak data underscore the significant association between listeriosis and diarrhea, with studies reporting symptom prevalence as high as 79% ^3,4^. The 2017–2018 South African outbreak, the largest recorded listeriosis event with over 1,000 cases and 216 deaths, underscores this risk. The outbreak went undetected for months until researchers identified a cluster of pediatric febrile gastroenteritis cases ^10^.

Clinical data suggest that inoculum dose critically affects diarrhea onset. Unlike hypervirulent enteric pathogens, such as *Shigella* and enterohemorrhagic *E. coli* (EHEC), which require less than100 cells, Lm-induced gastroenteritis typically requires high-dose exposure, often exceeding 10^10^ to 10^11^ CFU/mL ^3,15^. These outbreaks highlight the importance of gastroenteritis and prevention strategies for effective listeriosis management. Despite its clinical significance, the precise host-pathogen interaction underlying Lm-induced gastroenteritis remains poorly understood. This limited understanding has hindered early listeriosis diagnosis, leading to missed opportunities to intervene before infection progresses to the invasive phase ^7,8,10,11^.

While classical virulence factors like Internalin A (InlA) and listeriolysin O (LLO) are thought to contribute to tissue damage and gastroenteritis ^3,16,17^, their roles remain controversial. Many clinical Lm strains express truncated, non-functional InlA yet still cause listeriosis in humans ^18–20^ and guinea pigs ^21,22^. Furthermore, LLO secretion is significantly diminished in a low-oxygen intestinal lumen, raising uncertainty about its primary involvement in the enteric phase^23^.

In contrast, our previous work identified *Listeria* adhesion protein (LAP), a housekeeping moonlighting acetaldehyde alcohol dehydrogenase (AdhE), as a key virulence factor for Lm intestinal paracellular translocation ^24–27^. LAP interacts with the host epithelial receptor Hsp60 ^28–30^ to activate the NF-κB-MLCK (myosin light chain kinase) signaling axis, triggering the redistribution of junctional proteins such as claudin-1, occludin, and E-cadherin, and caveolar endocytosis to open paracellular gaps ^31,32^.

Additionally, physiological stressors such as hypoxia ^33,34^ and nutrient limitation ^35,36^ in the gut markedly upregulate LAP, reinforcing its crucial involvement in the early intestinal phase of infection.

In this study, we show that Lm-induced gastroenteritis occurs primarily after ingestion of a high dose of Lm, and gastroenteritis severity strongly correlates with LAP expression across Lm strains. We show that while diarrhea may act as a host defense mechanism to clear pathogens ^37^, local myeloid cell (CD11b^+^) hyperactivity, specifically neutrophils, drives disease severity. Furthermore, we show that LAP interacts with TLR2 and activates NF-κB in concert with LAP-Hsp60-mediated signaling pathways ^31^. TLR2-mediated innate activation is critical for protective immunity against Lm ^38^, as TLR2 deficiency abrogates protective immunity and reduces diarrheal symptoms in TLR2-deficient mice. Likewise, MLCK deficiency suppresses gastroenteritis. Our findings establish that, in addition to the LAP-TLR2 signaling pathway, MLCK activation is a critical driver of the immunopathology and ion channel dysfunction that characterize Lm-induced gastroenteritis at high infectious doses.

## Results

### LAP Mediates Epithelial Barrier Disruption and *Lm* Translocation by Widening the Paracellular Gap

Intestinal epithelial colonization is the crucial early step for subsequent invasion by enteric pathogens. Five gastroenteritis-causing Lm outbreak strains (1/2b and 4b serotypes) obtained from CDC showed strong adhesion (**Fig. 1a, Supplementary Table 1**), invasion (**Fig. 1b**), and paracellular translocation (**Fig. 1c**) across human intestinal enterocyte-like Caco-2 cell monolayers, while *lap*^─^ and Δ*inlA* showed reduced adhesion and invasion. As expected *lap*^─^ showed significantly impaired paracellular translocation, while Δ*inlA* showed translocation efficiency similar to WT (**Fig. 1c**). Immunoblot confirmed the expression of the Internalin family of proteins, such as InlA and InlB, on the bacterial cell wall and in the extracellular milieu (**Fig. 1d**). InlA expression is higher in serotype 1/2b than in 4b. In contrast, LAP was highly expressed in 4b serotypes compared with 1/2b serotypes (**Fig. 1d**). Together with our neuroinvasive F4244 (4b, CC6) strain ^39^, we chose H7014 (4b, CC2) for its greater adhesion, invasion, and paracellular translocation than other gastroenteritis-causing outbreak strains to support further studies. Interestingly, H7014 and F4244 share similar adhesion and translocation phenotypes (**Fig. 1a-c**) and comparative genomic features (**Supplementary Fig. 1a,b**).

**Figure 1.**
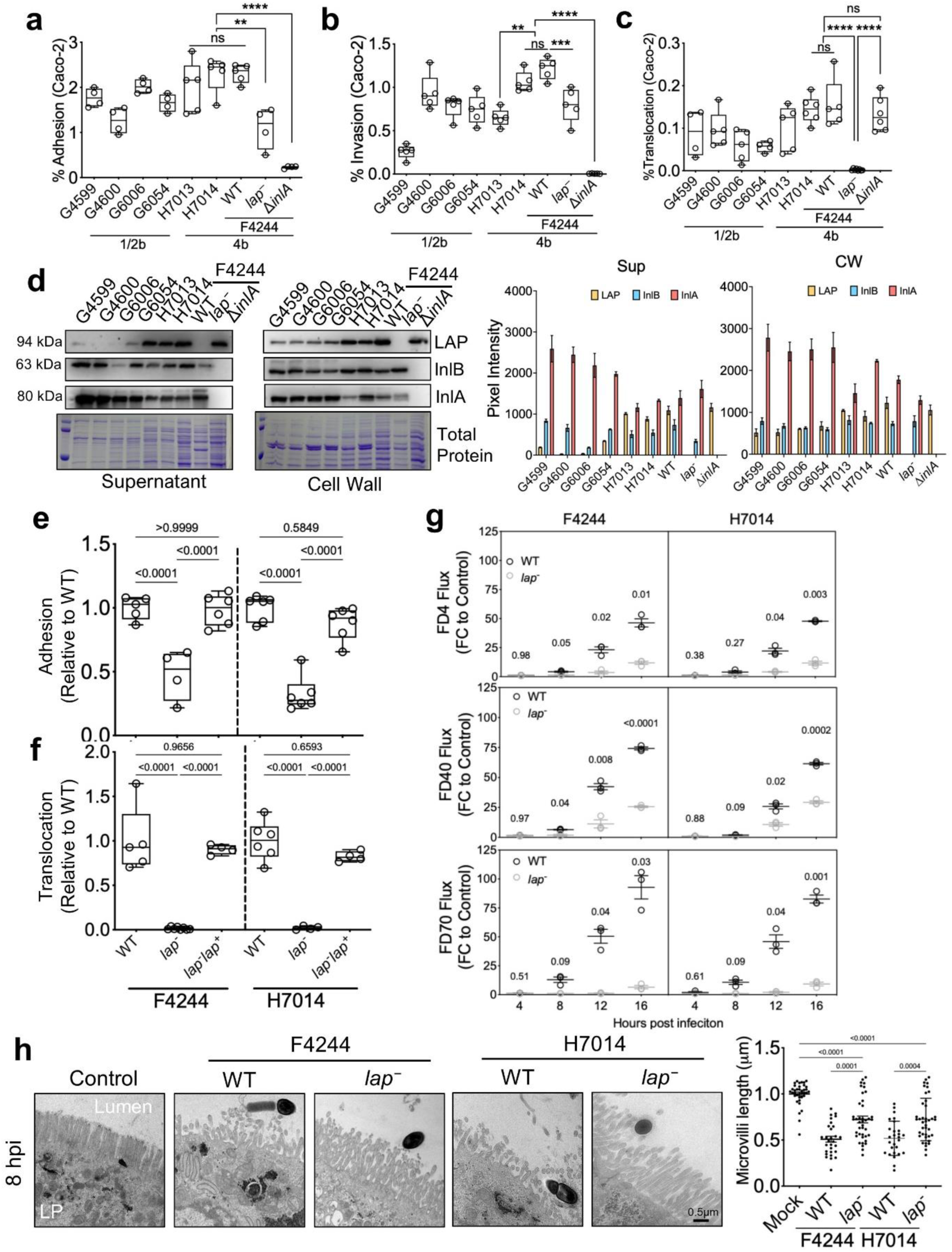
LAP Mediates Epithelial Barrier Disruption and Lm Translocation by Expanding Paracellular Gap. **a-c.** Quantification of adhesion (a), invasion (b), and translocation (c) of 7 gastroenteritis-causing Lm isolates (serotypes 1/2b and 4b) to the human intestinal Caco-2 cell line. **d.** Immunoblot analysis of LAP, InlA and InlB levels in Lm cell wall (CW) and extracellular supernatant (Sup). Coomassie blue staining of total protein (TP) is used as a loading control. Corresponding densitometry plots illustrate the relative expression of each protein across isolates. **e,f.** Comparison of adhesion (e) and translocation (f) of Lm WT, *lap*^-^ and *lap*^+^ complementary strains in both F4244 and H7014 backgrounds to Caco-2 cells. **g.** Time-dependent flux of FITC-dextran tracers (4, 40, 70 kDa) in Lm Caco-2 translocation assays. Plots show relative fold change in fluorescent intensity (FC) relative to the non-infected (gray) Caco-2 control. **h.** Representative scanning electron microscopy (TEM) images of Lm-infected Caco-2 at 8 hpi. The accompanying plot quantifies the average microvilli length following infection with Lm strains. All error bars represent TEM. ****p<0.0001; ***p<0.001; **p<0.01; *p<0.05; ns, no significance. All experiments were repeated at least three times unless otherwise noted.

To examine the LAP-mediated adhesion and translocation in the H7014 strain, we created an isogenic *lap-*deficient insertion mutant strain (*lap*^─^ H7014) (**Supplementary Fig. 1cd**), which also showed adhesion and paracellular translocation defects in human intestinal Caco-2 cells but was restored by *lap* complementation (*lap*^─^ *lap*^+^) (**Fig. 1e,f**). Additionally, whole-genome sequencing and analysis detected no significant differences in virulence-related genes between these two strains (**Supplementary Fig. 1a**). Further, Western blot analysis confirmed the absence of LAP in the *lap*^─^ H7014 strain while maintaining InlA and LLO expression (**Supplementary Fig. 1d**).

LAP can open the paracellular gap by disrupting the tight junctions ^31,32^, leading to diarrhea. However, the diameter of the gap opening was not known, so it was determined by monitoring the passage of varying molecular weight (MW) tracers: 4 (FD4), 40 (FD40), and 70 (FD70)-kDa dextran conjugated to fluorescein isothiocyanate (FITC) with a hydrodynamic pore diameter of 23 Å, 66 Å, and 103 Å, respectively ^40^. Time-dependent analysis of FD4, FD40 and FD70 movement through the epithelial monolayer revealed that exposure to H7014 and F4244 strains causes a ∼3, 2.5, and 7-fold increase in flux relative to corresponding isogenic *lap*^─^ strains at 4 h post-infection (hpi), which increased to more than 20-fold after 12 hpi (**Fig. 1g**). The significant decrease in TEER values in F4244 and H7014 WT but not in the *lap*^—^ counterpart, suggests a breach of the intestinal barrier (**Supplementary Fig. 1e**). Furthermore, transmission electron microscopy (TEM) of the Caco-2 monolayer at 8 hpi confirmed that infection with WT strains disrupted the absorptive epithelial brush border, showing decreased villus density and a damaged segmented-like structure. However, infection with their respective isogenic *lap*^─^ strains only slightly disrupted the cell monolayers (**Fig. 1h**). Overall, these results indicate that LAP plays a conserved role in Lm pathogenicity, contributing to bacterial adhesion and translocation across the epithelial barrier and inducing epithelial permeability.

### Acute Gastroenteritis Peaks at 48 hours Following High Infectious Dose

Outbreak reports indicate that Lm-induced diarrhea is associated with consuming heavily contaminated food, with inoculum levels reaching about 2.9 × 10^11^ CFU per person ^4^. To determine the bacterial load required to trigger gastroenteritis in a mouse model, C57BL/6 mice were orally administered three doses of Lm F4244 and H7014: 1 × 10^9^ (Low), 5 × 10^9^ (Medium), and 1-2 × 10^10^ (High) CFU/mouse.

At 48 hpi, the high-dose group (1-2 × 10^10^ CFU) showed severe intestinal inflammation, including fluid/gas accumulation and discoloration (yellowish to reddish) of both the small and large intestine (**Fig. 2a, Supplementary Fig. 2a**). Furthermore, high-dose infection led to progressive emptying of the ceca. Fecal pellets were softer and lighter in color than in lower-dose groups (**Fig. 2a, Supplementary Fig. 2a**). These clinical-pathological features were less severe at lower doses, demonstrating a dose-dependent relationship between Lm inoculum and gastroenteritis symptoms.

**Figure 2.**
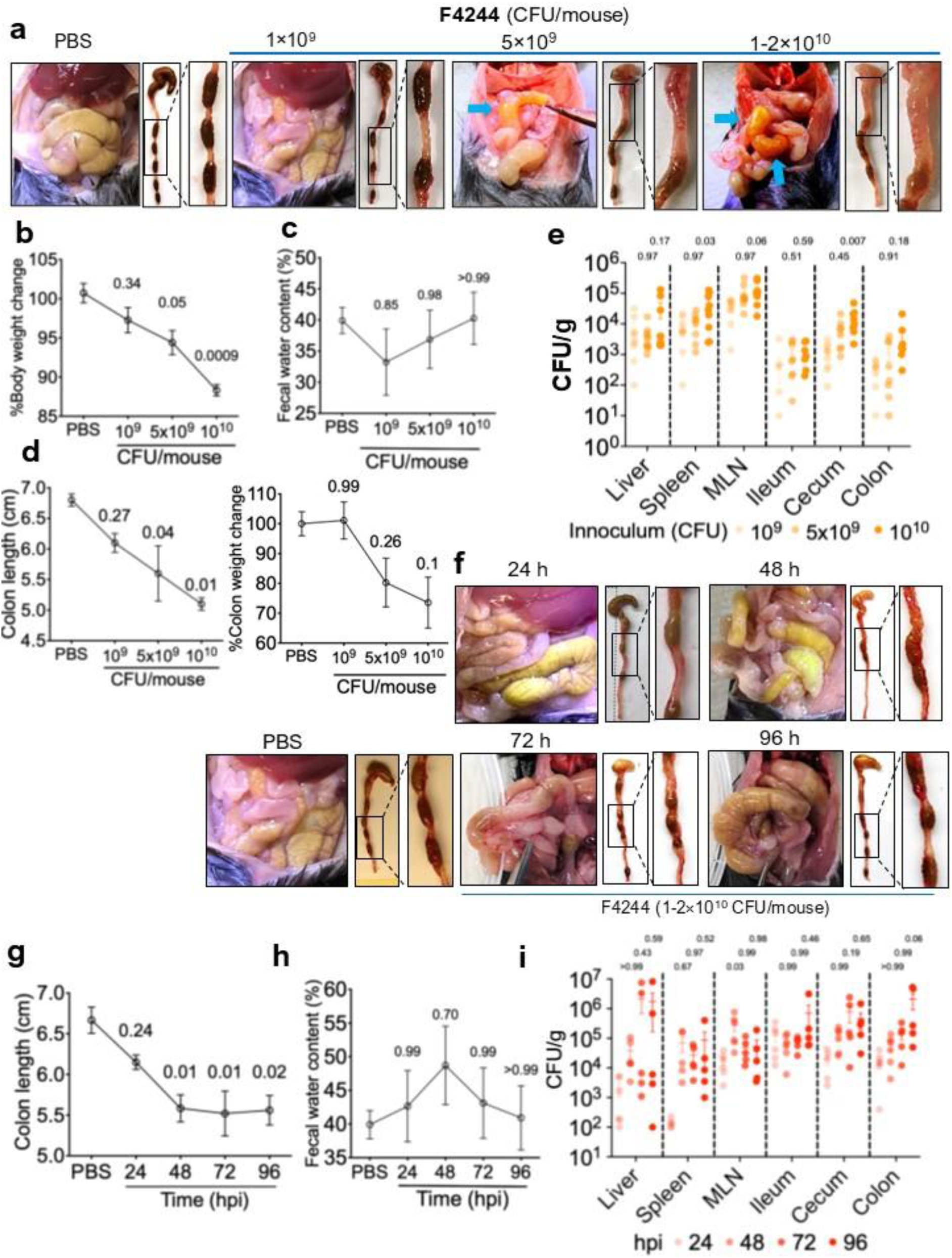
Acute Gastroenteritis Peaks at 48 hours Following High Infectious Dosea. Representative images of gross intestinal pathology of C57BL/6 WT mice that were intragastrically infected with 1 x 10^9^, 5 x 10^9^, 1-2 x 10^10^ CFU/mouse of Lm F4244, or PBS (Mock). **b.** Percentage change in total body weight and cecum-colon weight. n= 3 **c.** Analysis of fecal water content. PBS: n=6, Lm-infected mice: n=12-15 **d.** Measurement of colon length and % weight change after differential Lm dosing: n= 3 **e.** Quantification of bacterial burden in intestinal and extra-intestinal tissues after mice were infected with variable dosing of Lm F4244 48 hpi. **f.** Representative gross pathology of the abdominal region and large intestine pathology over time. **g.** Longitudinal measurement of colon length. n=5-6 **h.** Analysis of fecal water content. n=5-6, for 24h: Mock (PBS)=6, 24h=5, 48h=9, 96h=5. **i.** Temporal Lm tissue burden of mice at 24, 48, 72, and 96 hpi. n=5-6 All error bars represent SEM. All experiments were repeated at least three times unless otherwise noted.

This dose dependence was further reflected in progressive weight loss before sacrifice, increased fecal water content, and cecum and colon shrinkage. The group receiving the high dosage showed 11.68±0.72% total body weight loss, 26.45±8.57% colonic weight loss, 8.4±5.02% more fecal water retention, and 1.7±0.1 cm shorter colon length relative to the group that received PBS only (mock) (**Fig. 2b-d, Supplementary Fig. 2b-d**). Compared with the cohort receiving a low dose of inoculum, the high dose yielded significantly higher Lm burdens in the spleen, MLN, and cecum, while Lm burdens in the remaining tissues increased dose-dependently without reaching significance (**Fig. 2e**). These data indicate a high infectious dose is necessary to recapitulate gastroenteritis in a mouse model mimicking human epidemic cases ^3^.

The average incubation period for Lm-induced gastroenteritis in humans ranges from 24 to 72 h ^4,5,41^. Accordingly, we examined disease progression over 96 h following Lm inoculation with 1-2 × 10^10^ CFU/mouse. Interestingly, gastroenteritis symptoms, such as fluid accumulation and colon shortening, peaked at 48 hpi, and we didn’t observe worsening changes thereafter (**Fig. 2f,g**). Notably, fecal water retention also peaked at 48 hpi before gradually returning to baseline levels (**Fig. 2h**), even as bacterial burdens in intestinal and extraintestinal tissues continued to rise (**Fig. 2i**). These data suggest that the Lm-induced intestinal symptoms are transient. Consequently, we used a dose of 1-2 × 10^10^ CFU/mouse for 48 h for all subsequent experiments.

### LAP is the Primary Driver of Gastroenteritis In Vivo

Although LAP induces epithelial barrier dysfunction ^27,28,31,32^, we have not observed a specific role for LAP in gastroenteritis-associated diarrhea in our previous animal (mouse and gerbil) studies. This is because we used a low infectious dose (1 × 10^9^ CFU) in earlier trials ^31,32^, which was insufficient to induce diarrheal symptoms, as shown in our findings (**Fig. 2a-d, Supplementary Fig. 2a-e**). To investigate LAP’s role in gastroenteritis, we used our optimized gastroenteritis model, in which C57BL/6 (B6) mice were orally gavaged with Lm @1-2 × 10^10^ CFU/mouse.

At 48 hpi, mice receiving Lm WT strains (F4244 and H7014) showed comparable signs of gastroenteritis. In contrast, those receiving the *lap*^─^ mutant strains at the exact dosage showed no signs of gastroenteritis (**Fig. 3a**). The WT-infected animals revealed a swollen intestine (jejunum, ileum, cecum, and colon) and enlarged mesenteric lymph nodes, yellow mucoid ingesta, soft fecal pellets, and mucoid feces (**Supplementary Fig. 3a,b**). Fecal pellets collected from the colon after necropsy (F4244 and H7014; 1-2×10^10^) were soft and sticky and remained smeared near the mouth of the Eppendorf tube, while the pellets from PBS-treated mice were firm, non-sticky, and settled near the bottom of the Eppendorf tube during collection. The mock (PBS) or *lap*^─^ strain-infected animals showed natural dark fecal pellets with precise oval shapes (**Fig. 3b**). Fecal water content (**Fig. 3c**), diarrhea score (**Fig. 3d**), and serum FD40 (**Fig. 3e**) increased in Lm WT-infected mice while significantly decreasing in *lap*^─^ strain-infected mice. These observations were corroborated by significantly elevated bacterial burdens in the ileum, colon, MLN, liver, and spleen (**Fig. 3f,g**). Histopathology analysis revealed slightly elevated pathological scores and tissue damage in the ileum, and significantly elevated scores in the colon (**Supplementary Fig. 3c**).

**Figure 3.**
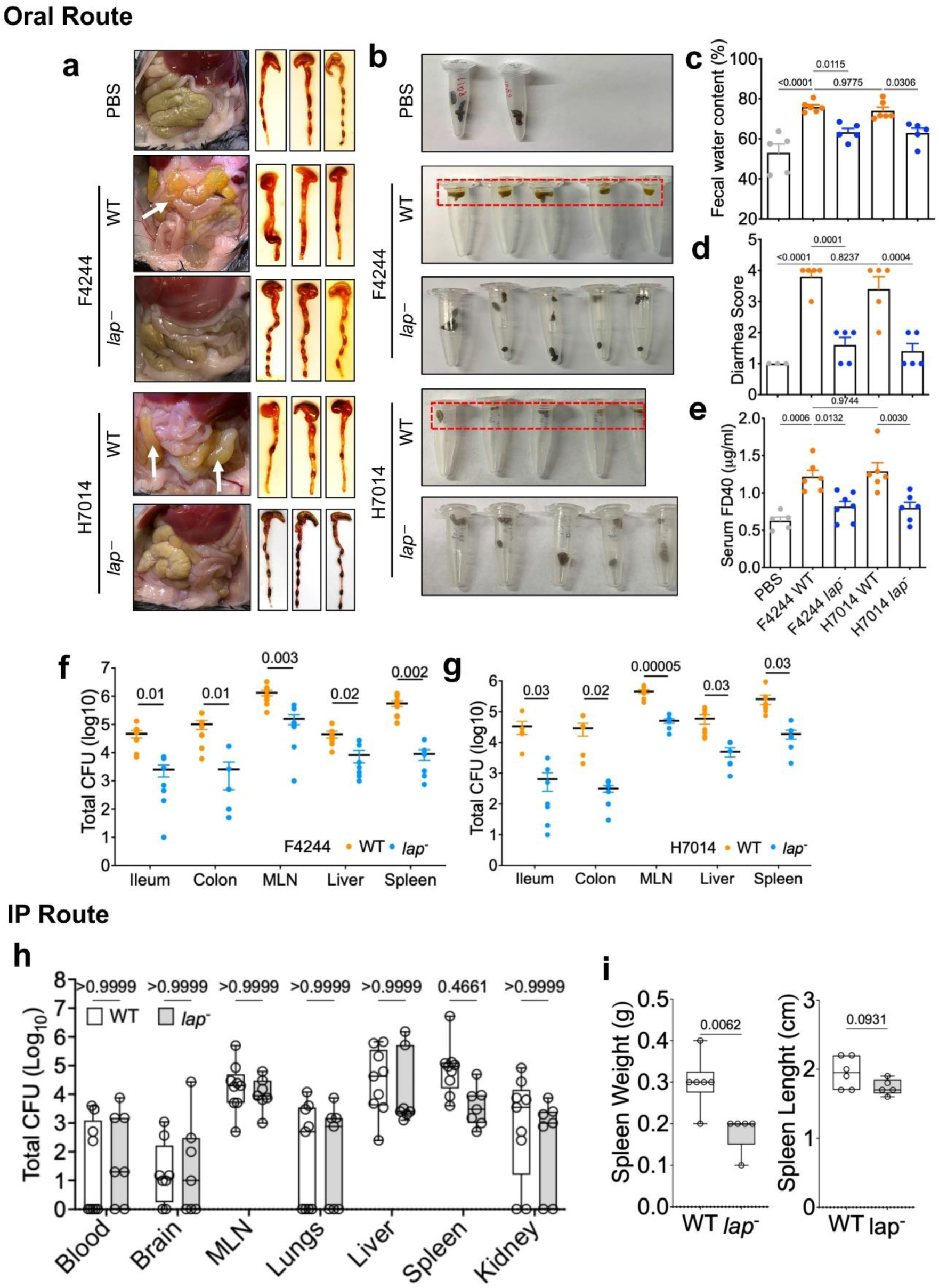
LAP is the Primary Driver of Gastroenteritis In Vivo. **a.** Representative gross intestinal pathology of wild-type B6 mice that were intragastrically challenged with 1-2 x 10^10^ CFU/mouse with WT or *lap^─^* strains of F4244 or H7014 at 48 hpi. **b.** Representative fecal morphology at 48 hpi; note the loss of pellet consistency in WT-infected mice compared to the firm, oval-shaped pellets in *lap^─^*-infected mice. **c-e.** Quantitative analysis of fecal water content (c), diarrhea scores (d), and intestinal permeability as measured by serum FD40 levels (e). **f,g.** Lm burden in intestinal and extra-intestinal tissues for F4244 (f) and H7014 (g) strains. **h.** Lm burden after i.p. injection in blood, brain, MLN, lungs, liver, spleen, and kidney after 96 hpi. **i.** Spleen weight and length All error bars represent SEM and are labeled with p-values. All experiments were repeated at least three times unless otherwise noted.

To determine whether the reduced intestinal and extra-intestinal burdens of *lap*^─^ reflect a specific defect in barrier crossing rather than a defect in *in vivo* survival and replication, we bypassed the gut barrier by inoculating mice intraperitoneally with WT or *lap*^─^ (5×10^6^ CFU/mouse) and enumerated bacteria in blood, brain, MLN, lung, liver, spleen, and kidney at 96 h post-infection. The two strains were recovered at comparable levels from each tissue/organ examined (**Fig. 3h**), consistent with our previous observation ^42^, indicating that *lap*^─^ is not attenuated for systemic survival, replication, or dissemination and that LAP contributes specifically to the intestinal phase of Lm translocation. Notably, despite equivalent bacterial burdens, both spleen weight and spleen length were significantly reduced in *lap*^─^-infected relative to WT-infected mice (**Fig. 3i**), suggesting LAP acts not only as an adhesin but as a direct immunostimulatory ligand.

Two other major virulence factors, LLO and InlA, could potentially contribute to gastroenteritis. All Lm strains were confirmed to be hemolytic on blood agar (**Supplementary Fig. 3d**). Interestingly, immunoblotting revealed consistent upregulation of LLO across both the *lap*^─^ strains (H7014 and F4244) (**Supplementary Fig. 1d**). The fact that *lap*^─^ strains elicited markedly milder gastroenteritis despite high LLO expression suggests that LLO may not be a primary driver of diarrheal symptoms, given the lesser adhesion of *lap*^─^ strains in this model. InlA involvement is harder to assess in mice, as wild-type InlA engages murine E-cadherin poorly ^43,44^, and InlA expression was unaltered in *lap*^─^ strains (**Supplementary Fig. 1d**). We therefore tested a murinized strain (*inlA^m^*; S192N/Y369S) ^34,45,46^ alongside the double mutant *inlA^m^ lap*^─^ (**Supplementary Fig. 3d,e**). Restoring InlA-E-cadherin engagement did not exacerbate diarrhea relative to WT. The Δ*inlA* retained a visible, albeit reduced, milder form of disease. In contrast, *inlA^m^ lap*^─^ was the most attenuated strain across fecal consistency, intestinal length, permeability, and tissue invasion (**Supplementary Fig. 3f-i**).

### LAP-induced Barrier Disruption Exacerbates Gastroenteritis Symptoms through Epithelial Cell Death and Immune Cell Infiltration

Both F4244 and H7014 strains produced similar gastroenteritis symptoms in the mouse model. Therefore, we used F4244 for all subsequent mechanistic experiments. In addition, we have several F4244-derived isogenic mutant strains that are also essential for unraveling mechanistic insights. At a low infectious dose, Lm translocates across the epithelial barrier and disseminates to deeper organs without replicating in the epithelial mucosa ^47^. However, in acute gastroenteritis, the intestinal epithelium is exposed to large amounts of Lm over a short interval, causing more severe damage and thus provoking substantially more intense inflammation.

As the intestine’s first barrier, intestinal epithelial cells (IEC) regulate the balance between fluid secretion and absorption ^48^. A disrupted epithelial surface is expected to perturb this balance. Flow cytometry analysis of murine intestinal cells shows that mice infected with Lm WT experienced increased IEC death in both the small and large intestine compared with the *lap^─^* strain. The small intestine is mainly affected, with more than half of the increased cell death (**Fig. 4a**). In addition, TEM of colonic tissue from Lm WT-infected mice shows shortened and disorganized microvilli, surface epithelial erosion, and loss of villus epithelial integrity relative to the *lap*^─^ or PBS control, where microvilli remained uniformly and compactly arrayed (**Fig. 4b**). These changes mirror the brush border disruption observed in the Caco-2 model (**Fig. 1h**). We also observed increased IL-1α (**Fig. 4c**), indicating substantial IEC lysis and immune activation ^49^.

**Figure 4.**
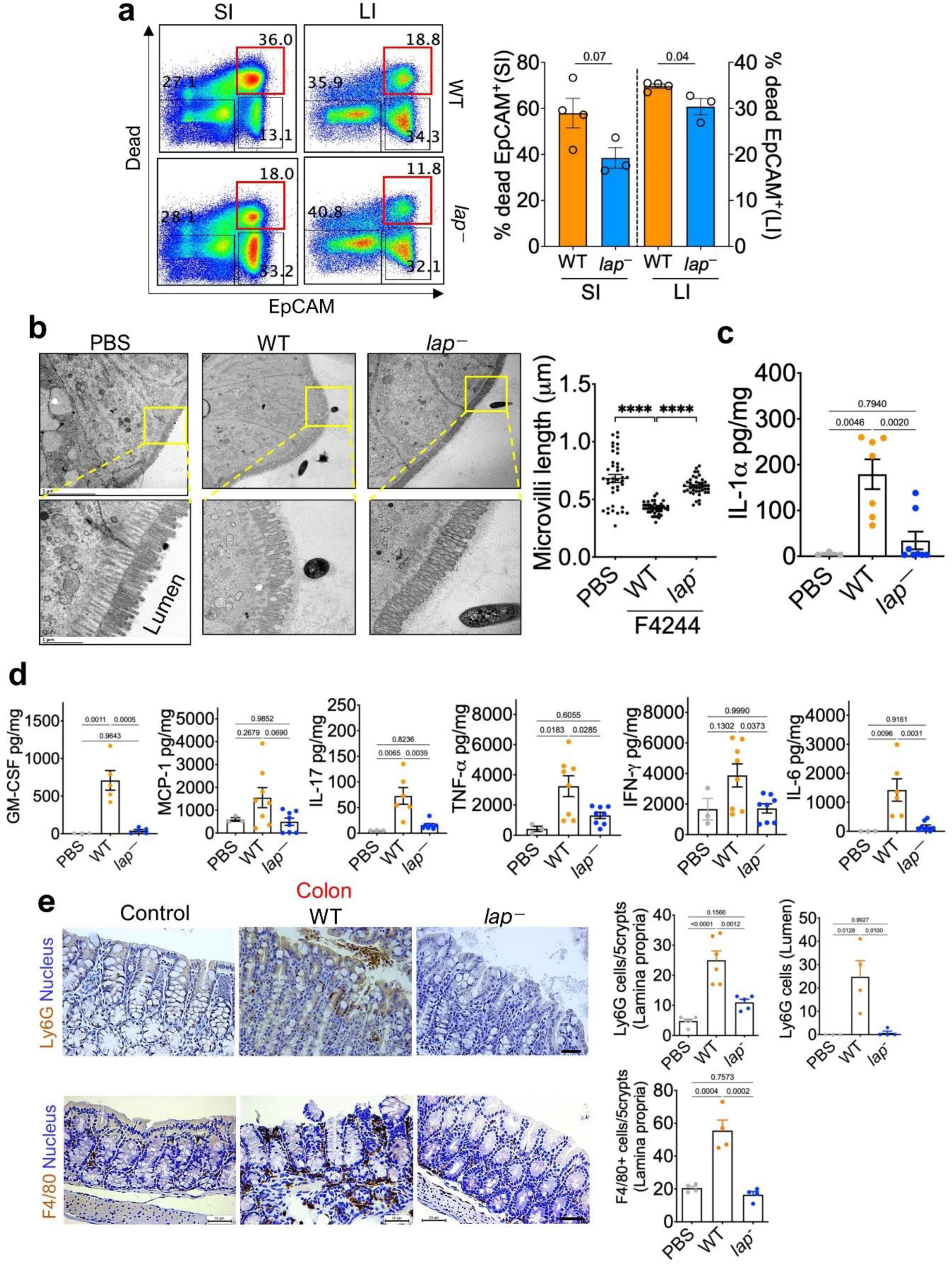
LAP-induced barrier disruption exacerbates gastroenteritis symptoms through epithelial cell death and immune cell infiltration. B6 mice were intragastrically challenged with 1-2 x 10^10^ CFU/mouse F4244 WT or *lap^─^* strains and sacrificed at 48 hpi. **a.** Flow cytometric analysis of IEC viability in the small intestine (SI) and large intestine (LI) comparing WT and *lap*⁻ infected mice — quantification of colonic cytokine levels. **b.** Representative TEM images of the colonic epithelial interface with morphology of microvilli. The plot shows the average microvilli length. **c.** Quantification of IL-1*α* Levels in the colon. n=5-6 **d.** Colonic proinflammatory cytokine profiling of mice infected with PBS, WT, and *lap*^―^. **e.** Representative IHC staining for Ly6G (brown) in the colon. Quantification of Ly6G+ neutrophil infiltration within the colonic lamina propria (average cells per 5 crypts per mouse) and the intestinal lumen (average cells per field per mouse). All error bars represent SEM. ****p<0.0001. All experiments were repeated at least three times unless otherwise noted.

The extent of inflammation is reflected in the colonic cytokine profile, with elevated pro-inflammatory cytokines (GM-CSF, MCP-1, IL-17, TNFα, IFNγ, IL-6) (**Fig. 4d**). During the initial phase of infection, the innate response is demonstrated by increased myeloid recruitment and activation (GM-CSF, MCP-1) and neutrophil-amplifying responses (IL-17, TNF-α), prompting us to investigate their infiltration into the intestine (**Fig. 4e, Supplementary Fig. 4a**). Immunohistochemistry of colonic sections revealed increased density of Ly6G⁺ neutrophils and F4/80⁺ macrophages in Lm WT-infected mice relative to *lap^─^*, distributed across the lamina propria and submucosa. Notably, Ly6G⁺ cells were also found in the colonic lumen in WT-treated mice, indicating transepithelial migration, whereas luminal neutrophils were rare in *lap^─^* infected mice (**Fig. 4e**). Ileal sections also showed enhanced infiltration of Ly6G⁺ cells (**Supplementary Fig. 4b**).

### CD11b⁺ Myeloid Cells Drive Diarrheal Pathology while Containing Lm Spread in the Small Intestine

Inflammatory myeloid cells are crucial innate defenders that protect against gastrointestinal pathogens, including Lm ^50^. However, the same effector programs that clear infection can also damage the epithelium. The resulting loss of absorptive surface and disruption of fluid-balance handling are recognized contributors to acute diarrhea. CD11b (Mac-1; *Itgam*) is the α-chain of the β2 integrin expressed on most innate myeloid cell lineages, including macrophages, neutrophils, and monocytes, and is critical for clearance of phagosome-coupled Lm ^51,52^.

To investigate the role of myeloid activation in Lm-induced gastroenteritis, wild-type C57BL/6 (B6) and *CD11b*⁻^/^⁻ mice were intragastrically infected with Lm at 1-2 × 10^10^ CFU/per mouse. At 48 hpi, gross pathology revealed severe intestinal inflammation in both groups, but with a striking distinction: the small intestine is severely inflamed in the B6 group, with shortened stomach-to-ileal segment and fluid accumulation, whereas these symptoms are relatively minor in *CD11b*⁻^/^⁻ mice (**Fig. 5a,b, Supplementary Fig. 7a**). Colon length was comparable between the two genotypes (**Supplementary Fig. 7a**). Although *CD11b*⁻^/^⁻ mice showed visibly reduced fecal content but both groups presented feces covered with yellowish sticky mucus (**Fig. 5c**).

**Figure 5.**
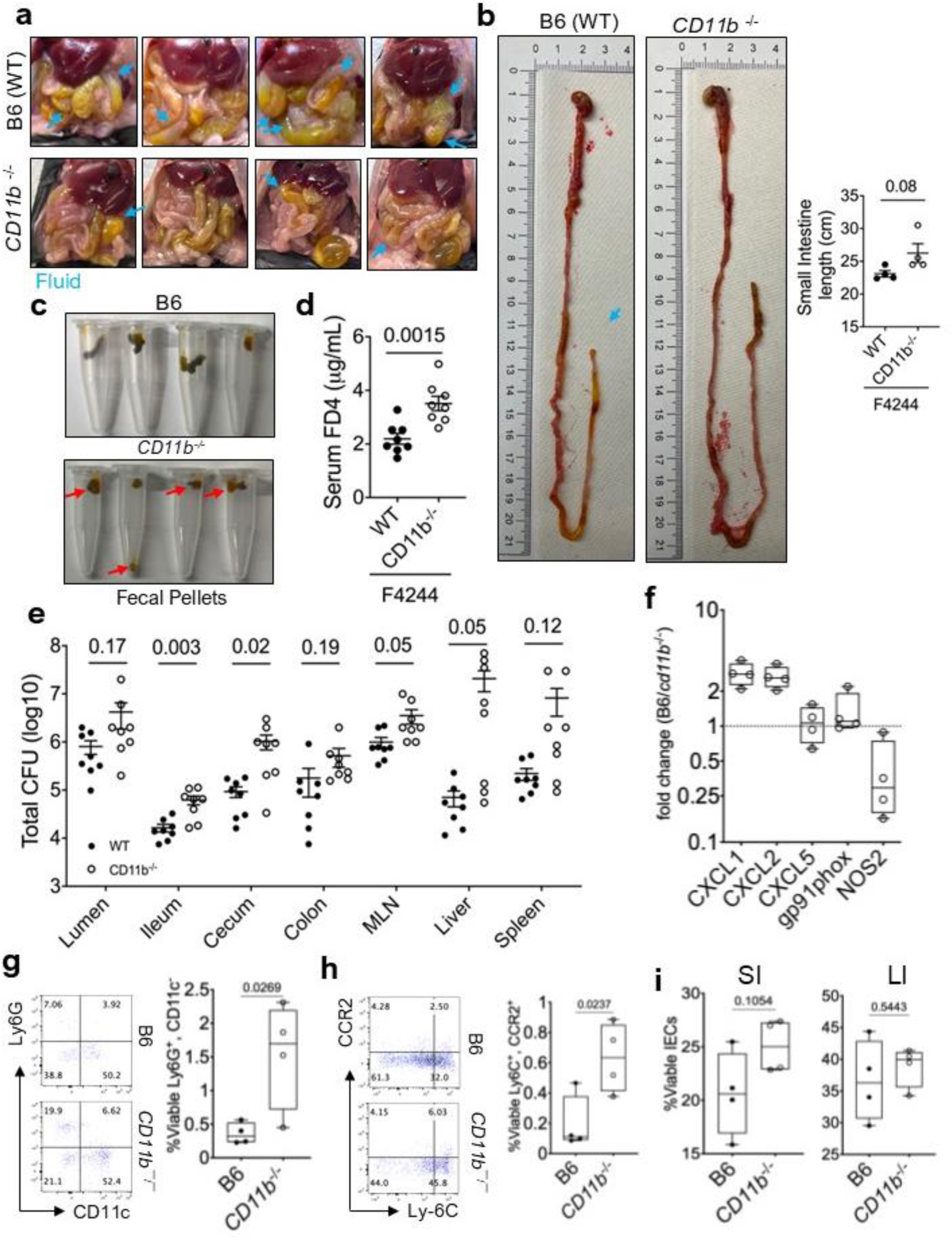
CD11b⁺ Myeloid Cells Mediate Diarrheal Symptoms that Contain Lm Spread in the Small Intestine. **a.** Representative images of the abdominal region of Lm-infected B6 (WT) and *cd11b*^⁻/⁻^ mice that were intragastrically challenged with 1-2 × 10^10^ CFU/mouse with Lm WT (F4244) at 48 hpi. Blue arrows indicate regions of fluid and mucus accumulation. **b.** Representative images and corresponding bar plots showing the length of the small intestine (stomach-ileum). **c.** Representative images of fecal morphology and volume collected at 48 hpi. Red arrows indicate the significantly reduced fecal volume and altered consistency in *CD11b*^⁻/⁻^ mice. **d.** Quantification of serum FD4 levels **e.** Lm burden in intestinal and extra-intestinal tissues at 48 hpi. **f.** Relative mRNA levels of inflammatory chemokines/cytokines from the ileum. **g. h.** Flow cytometric quantification of neutrophils (Ly6G^+^, CD11c^-^) (g) and inflammatory monocytes (Ly6C^+^, CCR2^+^) (h) from the small and large intestine. Representative density plots and viable cell counts are shown. **i.** Percentage of ileal viable IEC cell counts from Lm-infected B6 (WT) and *cd11b^⁻/⁻^* mice in ileum and colon. All error bars represent SEM labeled with p-values. All experiments were repeated at least three times unless otherwise noted.

This milder gross pathology in *CD11b*⁻^/^⁻ mice was unexpected, as they exhibited a more substantial breach of intestinal permeability, measured by increased serum FD4 levels (**Fig. 5d**). They also carried a markedly higher bacterial burden in both intestinal (ileum, cecum, colon) and extra-intestinal tissues (MLN, spleen, liver) (**Fig. 5e**). To assess myeloid function, we measured neutrophil chemokines and effector enzymes. CXCL1 and CXCL2 were higher in B6 than *CD11b*⁻/⁻ mice, whereas CXCL5 was unchanged, indicating acute myeloid recruitment rather than a global increase in epithelial chemoattractant output. cybb/gp91phox transcript slightly increased but not significant; NOS2 is modestly increased in *CD11b*⁻/⁻, which could be due to an expanded monocyte compartment and higher bacterial burden (**Fig. 5f**).

B6 mice also showed better myeloid recruitment, with 2-fold higher CXCL1, and CXCL2 transcript levels than *CD11b*⁻^/^⁻ (**Fig. 5f**). In addition, we measured the transcriptional levels of a set of genes that reflect the net fluid movement across the epithelium, including apical absorptive transporters *slc9a2*/NHE2, *slc9a3*/NHE3, *slc5a1*/SGLT1, *slc26a3*/DRA, the secretory chloride channel *cftr*, its basolateral partner *slc12a2*/NKCC1, and the calcium-activated chloride conductance regulator *clca1*. Interestingly, we found that B6 mice had both upregulated (SGLT1 and CFTR) and downregulated (DRA, NKCC1) transcripts for both absorptive and secretory ion channels or their regulator (CLCA1) ^53^ and compared to *CD11b*⁻^/^⁻ mice (**Supplementary Fig. 7b**), which might contribute to fluid imbalance in the gut.

Because *CD11b*⁻^/^⁻ mice are immunocompromised, we asked whether gastroenteritis symptoms in this genotype occurred earlier. However, at 24 or 48 hpi, we observed no signs of gastroenteritis (fluid accumulation, mucoid feces, and intestinal shortening) (**Supplementary Fig. 7c,d**). In parallel, we evaluated whether *lap*^─^ behaves differently in *CD11b*⁻^/^⁻ mice. The results show comparable gastroenteritis symptoms, fecal consistency, and bacterial burden in Lm WT and *lap*^─^ (**Supplementary Fig. 7c-e**), suggesting myeloid activation may contribute to LAP-mediated gastroenteritis.

This combination of high bacterial load and milder diarrheal symptoms in *CD11b*^⁻/⁻^ mice suggests that a dysfunctional innate immune response primarily drives diarrheal symptoms. In addition to neutrophils, CCR2^+^ monocytes are also crucial for controlling Lm infection ^54^. Characterization of the immune infiltrate revealed a significant increase in both Ly6G⁺CD11c⁻ neutrophils (**Fig. 5g**) and CCR2⁺Ly6C⁺ monocytes (**Fig. 5h**) in the small intestine of *CD11b*⁻^/^⁻ mice despite lower chemokine levels (**Fig. 5f)**. Mac-1 engagement is a pro-apoptotic signal in neutrophils; thus, CD11b-deficient neutrophils could show markedly delayed apoptosis, prolonging the lifespan (Coxon et al., 1996). Furthermore, IEC death is numerically higher in the small intestine than in the large intestine (**Fig. 5i, Supplementary Fig. 8 and 9**). Although we can’t establish a direct causal relationship between myeloid activation and Lm-induced gastroenteritis, reduced myeloid activation led to less IEC death in the small intestine, where fluid accumulation is most severe.

### TLR2 Signaling Mediates LAP-induced Gastroenteritis

To further characterize the innate signaling receptor for LAP, we performed gene set enrichment analysis (GSEA) on Lm-infected Caco-2 transcriptomes. The hallmark IL6-JAK-STAT3 signaling gene set was the top enriched pathway in Lm WT compared with *lap*^─^ (NES=1.55, *P* < 0.05), along with the leading-edge genes dominated by innate immune mediators including *Myd88*, *Tnfr*, *Il4r*, *Il6st*, *Tlr2*, etc in Caco-2 cell model (**Fig. 6a, Supplementary Fig. 10a-c**). Notably, among these genes, TLR2 is the only cell-surface pattern-recognition receptor that can directly transduce an extracellular bacterial protein signal and signal through MyD88 at the plasma membrane without a TRIF-dependent arm. These observations led us to hypothesize that LAP may directly engage TLR2 to activate NF-κB.

**Figure 6.**
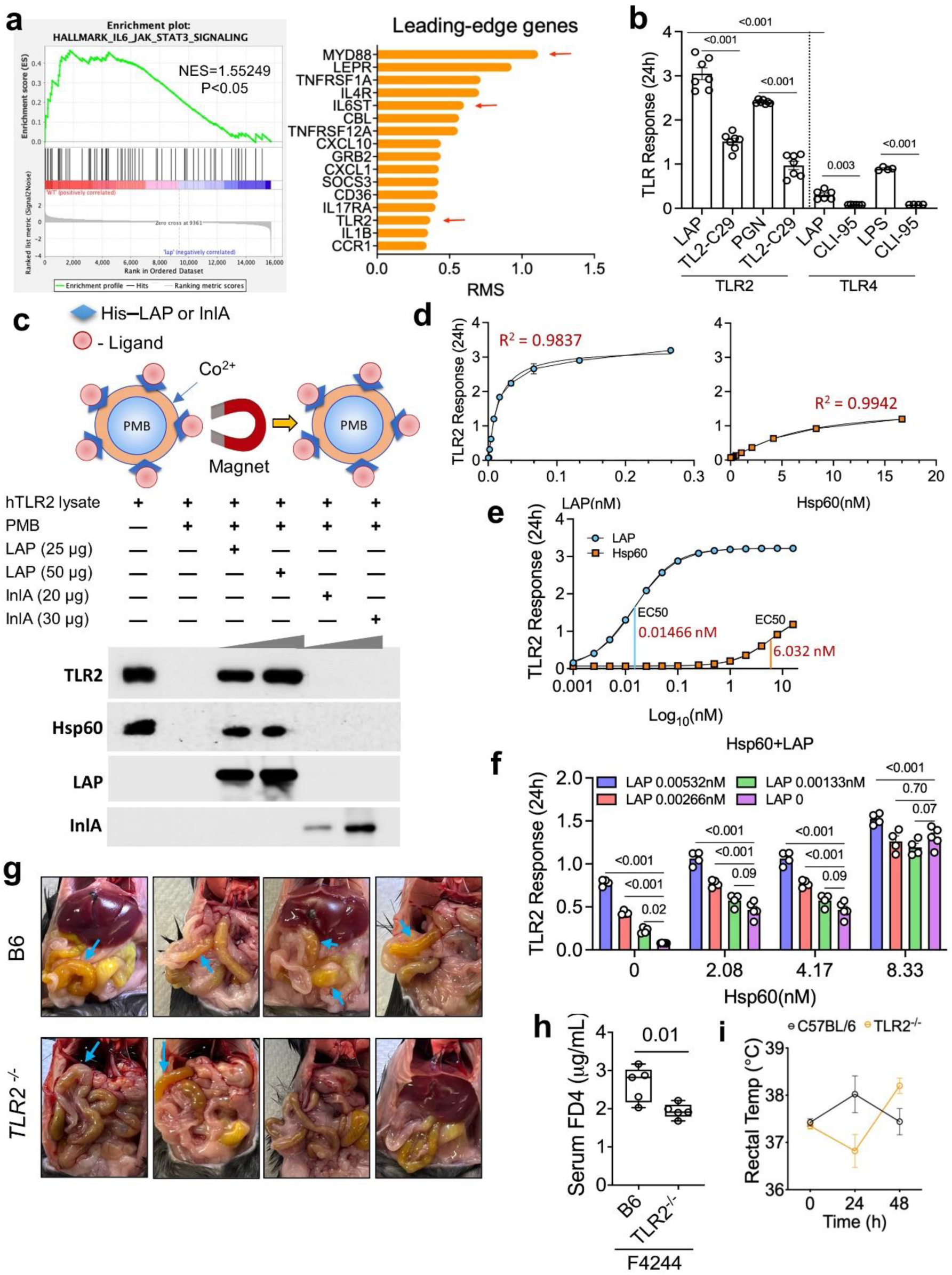
LAP-Induced Gastroenteritis is Mediated by TLR2 Signaling. **a.** Hallmark gene set enrichment analysis (GSEA) highlighting the activation of the IL6–JAK–STAT3 signaling pathways in Lm WT-infected Caco-2 cells over *lap*⁻. **b.** TLR response to purified LAP, PGN, LPS, and inhibitors (PGN inhibitor TL2-C29 and LPS inhibitor CLI-95) in HEK-Dual™ hTLR2 and HEK-Dual™ hTLR4 reporter cell lines after 24 h exposure. **c.** Pull-down assay for TLR2 from HEK-Dual™ hTLR2 cell lysate using paramagnetic beads (PMB) coated with LAP or InlA. **d.** TLR2 signal response curve (regression coefficient) for variable concentrations of LAP and Hsp60. **e.** Dose-response (EC_50_; half-maximal effective concentration) calculation for LAP and Hsp60. **f.** TLR2 response to LAP+Hsp60 used together at variable concentrations. **g.** Representative images of the intestinal morphology of Lm-infected C57BL/6 (B6) and *TLR2*^⁻/⁻^ mice at 48 hpi after intragastric challenge with 1-2 × 10^10^ CFU/mouse F4244 Lm WT. Blue arrows indicate regions of fluid and mucus accumulation. **h.** Assessment of systemic barrier breach as indicated by serum FD4 levels at 48 hpi. **i.** Rectal temperature recordings at 0, 24, and 48 hpi, illustrating the delayed febrile response in *TLR2*^⁻/⁻^ mice. All error bars represent SEM, with p-values. All experiments were repeated at least three times unless otherwise noted.

To test this hypothesis, we used the HEK-Dual™ hTLR2 (NF-κB-IL8) reporter cell line and compared the results with the TLR4 reporter cell line (HEK-Dual™ hTLR4 (NF-κB)). We observed that LAP-dependent NF-κB activation is TLR2-mediated and 7-fold more pronounced than TLR4 signaling (**Fig. 6b**), suggesting a dominant role for TLR2 signaling during LAP-TLR2 interaction. LAP also elicited a stronger response than the canonical TLR2 agonist peptidoglycan (PGN) (**Fig. 6b**). Treatment with the TLR2 inhibitor TL2-C29 significantly attenuated the LAP-induced signaling while maintaining cell vaibality (>80%; **Supplementary Fig. 11a**). In contrast, InlA used as a control, did not stimulate TLR2 signaling (**Supplementary Fig. 11b**). To verify LAP interaction with TLR2 further, we performed a pulldown assay where LAP coated paramagnetic bead (PMB) recovered TLR2 from the cell lysate obtained from the HEK-Dual™ hTLR2 reporter cell line (**Fig. 6c**), supporting association between LAP and TLR2. InlA, used as a control, did not interact with TLR2 in the cell lysate. As expected, LAP also recovered Hsp60 from the cell lysate (**Fig. 6c**), consistent with our previous observation ^28,29^, and suggesting that LAP-Hsp60-mediated NF-κB signaling ^31,32^ is active during gastroenteritis.

Furthermore, LAP-mediated TLR2 activation is dose-dependent (**Fig. 6d**); likewise, Hsp60, a TLR2 ligand ^55^, also initiated TLR2 signaling in a dose-dependent manner (**Fig. 6e**). Nonlinear regression yielded EC50 (half-maximal effective concentration) values of approximately 0.015 nM for LAP and 6.02 nM for Hsp60, indicating that LAP was approximately about 400-fold more potent than Hsp60 on a molar basis (**Fig. 6e; Supplementary Fig. 11c**). Combined LAP and Hsp60 treatment produced a concentration-dependent TLR2 response. Two-way ANOVA revealed significant effects of both LAP and Hsp60 at variable concentrations, as well as a significant LAP×Hsp60 interaction (*P* < 0.001). Consistent with this interaction, the fold increase induced by Hsp60 relative to the corresponding LAP-alone baseline progressively decreased as LAP concentration increased. Although both LAP and Hsp60 stimulated TLR2, Hsp60’s incremental contribution diminished as LAP concentration increased, suggesting that additional stimulation became progressively smaller at higher combined concentrations (**Fig. 6f**).

We also assessed TLR2 activation by live Lm F4244 WT and *lap^─^*cells; however, at 6 h, they did not show any apparent differences, whereas at 24 h, the TLR2 response in WT was significantly higher than that in the *lap^─^* strain (**Supplementary Fig. 11d**). However, both WT and *lap^─^* treated cells (MOI 10 or 1) showed significant hTLR2 HEK cell death at 24 h, indicating that TLR2 activation requires an optimal time and bacteria-to-HEK cell ratio. Furthermore, PGN analysis in WT and *lap^─^*did not reveal significant differences (**Supplementary Fig. 11e**). Collectively, these data show that LAP activates TLR2 signaling for canonical NF-κB activation, highlighting the importance of the LAP-TLR2 axis in Lm pathogenesis.

We next asked whether this LAP-TLR2 axis drives intestinal pathology in vivo using TLR2-deficient (*TLR2*⁻^/^⁻) mice. *TLR2*⁻^/^⁻ and B6 (WT) mice were intragastrically gavaged with a gastroenteritis-inducing dose of Lm (1-2×10^10^ CFU/mouse). At 48 hpi, *TLR2*⁻^/^⁻ mice exhibited a striking alleviation of gastroenteritis symptoms compared with B6 mice. This was evidenced by significantly attenuated fluid accumulation (**Fig. 6g**), reduced intestinal permeability (lower serum FD4 levels; **Fig. 6h**), and a delayed febrile response (**Fig. 6i**) compared to B6. Moreover, significant shortening of the small intestine was observed in B6 mice, whereas *TLR2*⁻^/^⁻ mice maintained normal tissue length (**Extended Fig. 11f**). Although colon length was comparable between the two groups (**Extended Fig. 11g**), *TLR2*⁻^/^⁻ mice had significantly diminished fecal content (**Extended Fig. 11h**), consistent with reduced diarrheal fluid volume. Strikingly, this profound reduction in gastrointestinal pathology occurred despite a bacterial burden comparable to that of B6 (**Extended Fig. 11i**).

We further tested the effect of Lm WT and *lap^─^* strain in *TLR2*⁻^/^⁻ mice following intragastric gavage. Interestingly, we observed no differences between the two strains in small intestinal length, cecum-colon length, FD4 flux, or bacterial tissue burden in intestinal (except the cecum) and extra-intestinal tissues (**Supplementary Fig. 12a-d**), suggesting the LAP-TLR2 axis is crucial for intestinal inflammation.

Expression of the Na⁺/H⁺ exchangers *Slc9a3/*NHE3 and *Slc9a2/*NHE2 also differed between genotypes, indicating TLR2-dependent transcriptional changes in ion transport. Regarding inflammatory markers, both acute and latent neutrophil- and macrophage-recruiting chemokines (*CXCL1*, *CXCL2*, *CXCL5*) were elevated in B6 mice compared with *TLR2*⁻^/^⁻ mice (**Supplementary Fig. 12e**). In contrast, the ROS-generating enzyme *Cybb*/*gp91phox*) was higher in *TLR2*⁻^/^⁻, and NOS2 levels were comparable between groups. Collectively, these data suggest that Lm-induced gastroenteritis (fluid accumulation, tissue shortening, and permeability loss) is driven by a TLR2-dependent host inflammatory response.

### MLCK Deficiency Preserves the Epithelial Barrier Integrity and Attenuates LAP-induced Gastroenteritis

The long isoform of myosin light chain kinase (long MLCK, MLCK 210) is a well-established regulator of tight junction (TJ) permeability and a primary driver of diarrheal pathology and “leaky gut” syndromes ^56,57^. We previously showed that LAP activates canonical NF-κB signaling and MLCK, redistributing claudin-1, occludin, and E-cadherin to permit paracellular low-dose Lm translocation 1 × 10⁹ CFU/mouse, and that this translocation is attenuated in long MLCK⁻^/^⁻ (referred to as *MLCK*⁻^/^⁻ later in the text) mice. However, that inoculum produces translocation and systemic dissemination without overt gastroenteritis. Because the junctional space opened by MLCK also serves as the route through which neutrophils complete transepithelial migration, we asked whether long MLCK is required not only for bacterial entry but also for the myeloid-driven pathology of gastroenteritis.

When mice received oral gavage of Lm @1-2 × 10^10^ CFU/mouse, we observed that *MLCK*⁻^/^⁻ mice were markedly resistant to Lm infection, with attenuated gastroenteritis symptoms and lower serum FD40 flux compared to WT B6 mice (**Fig. 7a,b**). MLCK deficiency also eliminated the LAP-dependent bacterial burden at this dose (**Fig. 7c**). Lm WT counts in intestinal and extra-intestinal tissues fell to the level of the *lap^─^* strain, whereas *lap^─^* burdens were unchanged from those in B6 mice.

**Figure 7.**
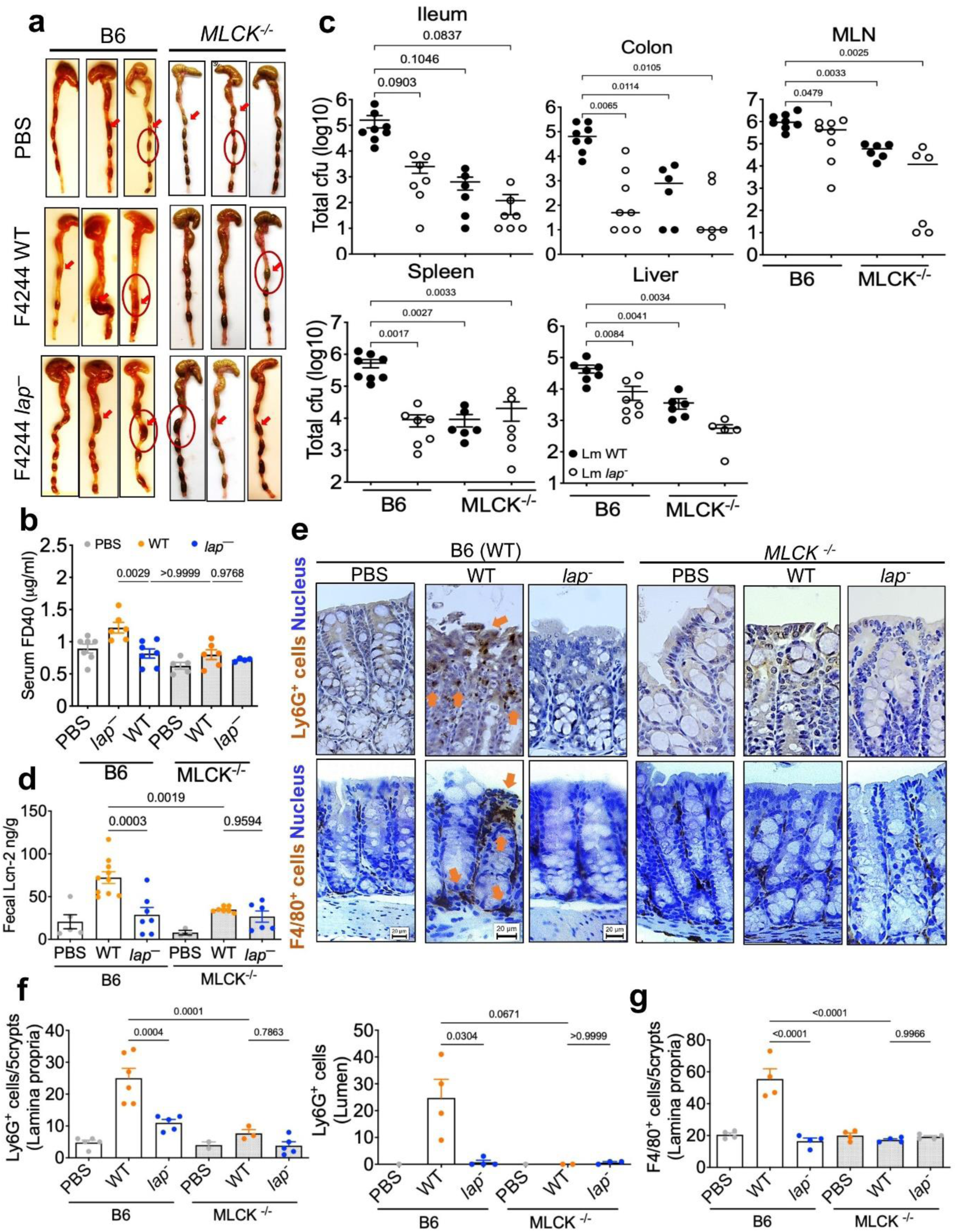
MLCK deficiency preserves the paracellular gap and attenuates LAP-induced gastroenteritis. **a.** Representative large intestinal (cecum-colon) pathology from B6 and *MLCK^⁻/⁻^* mice at 48 hpi following Lm intragastric gavage. Red arrows indicate regions of fluid and mucus accumulation. **b.** Gut permeability assessment using serum FD40 levels. **c.** Lm burden in intestinal and extraintestinal tissues. **d.** Fecal lipocalin (Lcn-2) levels at 48 hpi. **e.** Representative colonic IHC staining of Ly6G^+^ (neutrophils) and F4/80^+^ (macrophages) shows enhanced recruitment to the lamina propria and lumen tissues from the Lm WT-treated group. The nucleus was counterstained with hematoxylin. **f,g.** Quantitative analysis showing the mean counts of Ly6G⁺ (f) cells in the lumen and lamina propria and F4/80⁺ (g) cells in the lamina propria. All error bars represent SEM and are labeled with p-values. All experiments were repeated at least three times unless otherwise noted.

In addition, *MLCK⁻^/^⁻* mice exhibited significantly lower levels of intestinal inflammation, as indicated by reduced fecal lipocalin-2 (LCN-2) levels (**Fig. 7d**) and intestinal histopathology analysis (**Supplementary Fig. 13a,b**). Furthermore, the characteristic increase in neutrophils and macrophage infiltration observed during Lm WT infection was absent in *MLCK⁻^/^⁻* mice (**Fig. 7e,f**). *MLCK⁻^/^⁻* mice showed fewer cleaved caspase-3⁺ cells within the villus extrusion zone and a smaller crypt Ki67⁺ compartment than B6 mice (**Supplementary Fig. 14a,b**), indicating attenuated epithelial turnover. Intestinal (ileum and colon) immunofluorescent staining revealed that *MLCK⁻^/^⁻* mice preserved expression of TJ and adherens junction (AJ)-associated proteins, including claudin-1, occludin, ZO-1, and E-cadherin, regardless of being challenged with WT or *lap^─^* strains (**Supplementary Fig. 15a,b**).

## Discussion

*Listeria monocytogenes*-induced gastroenteritis is often underrecognized because it is transient and self-limiting. However, identifying this enteric phase is clinically important for preventing progression to life-threatening systemic listeriosis. The nonspecific nature of these symptoms makes it diagnostically challenging to link them specifically to Lm ^2,3,5^. To bridge this gap, we used murine models to show that Lm-induced gastroenteritis is not merely a passive result of bacterial colonization, but an active, LAP-dependent immunopathological response driven by the host’s innate immune system.

The capacity to induce gastroenteritis was not restricted to a single lineage or clinical isolate. Both the gastroenteritis strain (H7014, CC2) and the neuroinvasive strain (F4244, CC6) induced gastroenteritis, despite differences in clinical origins and genetic backgrounds ^39^ (**Supplementary Fig. 1a; Supplementary Table 1**). F4244 belongs to Clonal Complex 6 (CC6), a lineage that frequently carries virulence islands such as LIPI-4 that enhance systemic potency ^58,59^. Across the 6 strains we tested, disease severity tracked with infectivity and LAP expression level. This is consistent with LAP-dependent pathology, although the limited number of strains examined precludes a formal correlation.

Severe gastroenteritis in mice required a high bacterial inoculum (1-2 × 10^10^ CFU), consistent with human outbreak reports estimating infectious doses between 10^10^ - 10^11^ CFU ^3,5,60^. Gastroenteritis peaked at 48 hpi and subsequently resolved, mirroring the clinical course of human febrile listerial gastroenteritis.

A central conclusion of this study is that the classical virulence factors InlA and LLO play minor roles in inducing Lm gastroenteritis. Although wild-type InlA is essential for epithelial entry (transcytosis)^61^ in humans, it plays a diminished role in mice due to species-specific incompatibility with E-cadherin ^43^. InlA^m^ strains, which restore E-cadherin-dependent epithelial invasion in mice, did not exacerbate diarrhea (**Supplementary Fig. 3f-i**), suggesting that enhanced epithelial invasion is insufficient to drive gastroenteritis pathology. In contrast, *lap*^─^ strain retaining functional LLO failed to induce symptoms (**Supplementary Fig. 1d; Fig. 3**), indicating that LLO is neither sufficient for nor the primary initiator of the inflammatory cascade driving fluid loss ^62^. These findings help clarify why LLO-based serology has limited diagnostic value for gastroenteritis ^63^.

Mechanistically, our data identify LAP-dependent activation of the TLR2-NF-κB axis as the molecular switch initiating this response at high infectious dose. Epithelial TLR2 expression extends from the small intestine to the proximal colon ^64^ and serves as a key surveillance mechanism for intestinal pathogens. During Lm infection, epithelial injury and myeloid activation are coupled, amplifying gut inflammation from the small to the proximal large intestine. IL-1α, an alarmin released from damaged epithelium, was elevated in Lm WT-infected mice, linking barrier disruption to downstream myeloid recruitment and activation ^65^. IL-1α and other proinflammatory cytokine/chemokines promotes neutrophil recruitment/activation across mucosal and other tissues but can also exacerbate collateral tissue damage ^66–69^. This pathology mirrors other inflammatory bowel conditions in which neutrophil activity correlates with epithelial injury ^65^.

*CD11b⁻^/^⁻* mice provide causal support for this interpretation. Despite carrying higher Lm burdens in the ileum, cecum, mesenteric lymph node, spleen, and liver, these mice displayed attenuated diarrheal symptoms (**Fig. 5a)**. Because luminal burden was comparable between genotypes, while burden rose in every tissue compartment, the difference reflects impaired containment of bacteria that have crossed the epithelium faster. Mucosal myeloid frequencies were also comparable between WT (B6) and *CD11b⁻^/^⁻* mice, indicating that the attenuated phenotype does not reflect a failed recruitment. Maintained cell numbers despite lower *CXCL1* and *CXCL2* are consistent with the prolonged survival of CD11b-deficient myeloid cells ^70,71^. Gastroenteritis severity therefore aligns with myeloid effector function rather than with bacterial load or myeloid abundance. The combination of increased systemic dissemination with muted enteric symptoms in *CD11b⁻^/^⁻* mice recapitulates the clinical picture of immunocompromised patients where gastrointestinal symptoms are mild or absent even as infection progresses to systemic listeriosis.

Lymphoid population profiling in the small intestine showed a modest shift, with a small increase in CD4⁺ and CD8⁺ cells in *lap^─^* animals and an increase in CD4⁻CD8⁻ double-negative cells in the Lm WT group. Myeloid population frequencies did not change significantly (**Supplementary Fig. 5 and 6**), indicating that the acute myeloid response is regionally concentrated rather than a global expansion of the intestinal immune compartment.

Downstream of TLR2-NF-κB-driven MLCK activation, myosin light chain (MLC) phosphorylation prompts tight junction opening ^56^. CD11b is essential for effective myeloid cell activation via TLR co-signaling ^72^. Because tight junctions gate paracellular immune cell migration ^57,73^, this creates the paracellular gaps necessary for neutrophil transmigration ^74^. Consistent with this mechanism, we observed elevated fecal lipocalin-2 (**Fig. 7d**), a marker of neutrophil degranulation ^75–77^, in WT infected mice relative to *lap^─^*, indicating that LAP-dependent junctional opening is accompanied by active neutrophil recruitment/activation at the mucosal surface. The chemokine component of this response was regionally patterned. Furthermore, contributions of the LAP-Hsp60-MLCK axis to epithelial tight junction opening for Lm transmigration ^31,32^, and possibly neutrophil passage through paracellular gaps cannot be ruled out; nevertheless, at high infectious doses (1-2×10^10^ CFU/mouse), the LAP-TLR2 axis coupled with MLCK activation appeared dominant during gastroenteritis onset.

The resulting fluid loss is malabsorptive rather than secretory because the functional absorptive surface is lost. At the ultrastructural level, LAP-dependent shortening of brush border microvilli in Caco-2 monolayers directly reduces absorptive membrane area. At the organ level, villus length was also reduced in Lm WT-infected mice relative to *lap^─^* infected mice, reproducing this loss in vivo. Consistent with these findings, CC3 was elevated at the villus tip, and Ki67 was elevated in the crypt of Lm WT-infected mice, indicating accelerated epithelial turnover with compensatory proliferation: mature, terminally differentiated absorptive enterocytes are lost from the villus tip and replaced by immature, crypt-derived cells with lower absorptive capacity. Secretory channels were not uniformly upregulated, and *CFTR* was reduced in Lm WT relative to *lap*^─^-infected mice (**Supplementary Fig. 4c**), arguing against a secretory mechanism of fluid loss.

Several limitations of this work should be noted. Both *TLR2⁻^/^⁻* and *CD11b⁻^/^⁻* are germline knockouts, so epithelial and myeloid contributions to TLR2 signaling cannot be separated in the current models, and developmental compensation cannot be excluded. A conditional, lineage-restricted deletion would resolve which TLR2 compartment initiates the responses. Because LAP partially engages TLR2 signaling, alternative receptor(s) may mediate LAP-MyD88 signaling, including IL-1R1, IL-18R, and ST2.

In conclusion, our findings identify LAP as a central mediator of Lm-induced gastroenteritis (**Fig. 8**). At high infectious doses, LAP recruits proinflammatory myeloid cells through the TLR2-MLCK signaling axis, triggering a host response that controls bacterial spread but also drives immunopathology and loss of absorptive surface, culminating in diarrhea. Furthermore, Hsp60, which interacts with both TLR2 and LAP, cannot be overlooked, as it may contribute to the onset of diarrheal symptoms and warrants further investigation. Nevertheless, our findings redefine *L. monocytogenes*-induced gastroenteritis as a host-driven inflammatory disease initiated by bacterial adhesion and tight junction opening rather than a direct consequence of bacterial invasion. Critically, symptomatic gastroenteritis reflects a competent myeloid response and an early, manageable stage of infection, and recognizing it offers a window for intervention before systemic dissemination.

**Figure 8.**
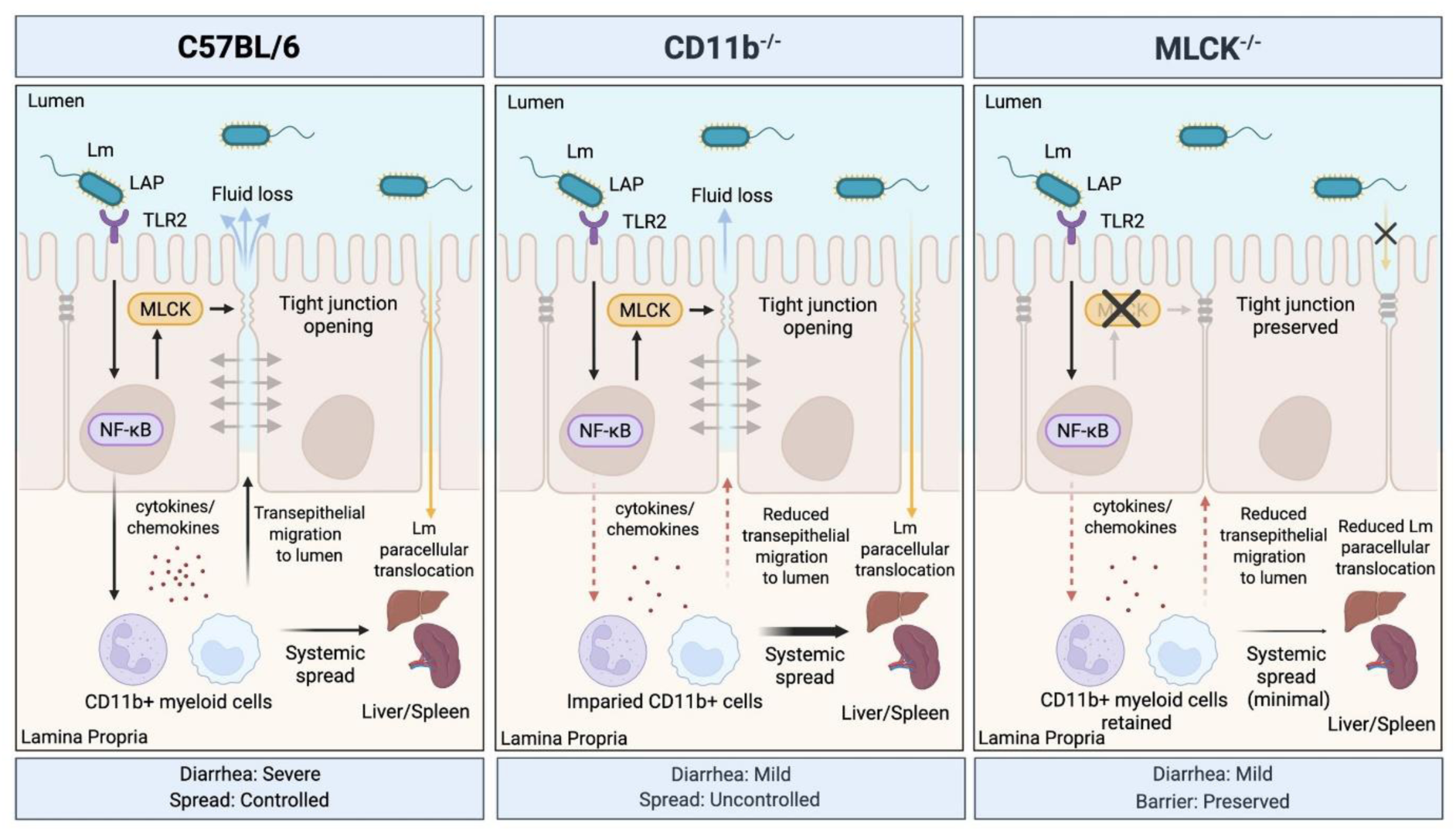
Summary of LAP-TLR2-NF-κB-MLCK signaling cascade in *L. monocytogenes*-mediated gastroenteritis.

## Acknowledgements

This work was supported by the U.S. Department of Agriculture National Institute of Food and Agriculture (Hatch accession no. 1016249) and the Bilsland Fellowship to D. Liu. Any opinions, findings, conclusions, or recommendations expressed in this publication are those of the authors and do not necessarily reflect the views of the USDA. We acknowledge the technical assistance of M. Samaddar, C. Li, and Peter Garner-Schmidt (CDC) in providing Lm outbreak strains.

## Author Contributions

DL, ST, AKB designed and performed the experiments, interpreted the data, and wrote the manuscript. DL, AKB, ST, XB, NG, LX, NIT, VN, YF, ZG, and RD performed mouse necropsy and bacterial burden determination. DL, JYF, SP, MRO designed and analyzed the flow cytometry. ST performed TEM and confocal imaging. AC assisted with IHC staining and histopathological analysis. AKB, MRO, AC, and LR critically edited the manuscript. All authors reviewed the manuscript.

## Resource Availability

All Lm clones are available upon request. Genetically modified mice are available from Jackson Lab. MLCK KO mice were supplied by J.R. Turner (Harvard Medical School). Processed RNA sequencing data is uploaded on NCBI database.

## Methods

### Bacterial Strains and Growth Conditions

All bacterial strains used are listed in **Supplementary Tables 1-3**, including Lm F4244 wild type (WT F4244, serotype 4b, isolated from cerebrospinal fluid of a patient) ^39^, isogenic *lap*-deficient insertion mutant (*lap*^─^F4244), *lap-*complemented (*lap*^─^ *lap*^+^F4244) ^24^, murinized InlA (mInlA), *lap*-deficient insertion mutant in mInlA (mInlA *lap*^─^)^32^, *ΔinlA* F4244 in-frame deletion mutant ^28^, 10403S in-frame deletion mutant *Δhly* (DP-L2161), human gastroenteritis causing Lm wild type strains (G4600, G6006, G6054, H7013 and H7014) and isogenic *lap*-deficient insertion mutant (*lap*^─^H7014). All *Lm* strains were grown in Tryptic Soy Broth supplemented with 0.6% yeast extract (TSBYE) for 14-16 h with shaking at 150 rpm at 37°C, unless otherwise stated. The *lap*^─^, *lap*^─^ *lap*^+^, and mInlA *lap*^─^ strains were grown in TSBYE supplemented with erythromycin or chloramphenicol, or both at 42°C. Detailed corresponding culturing conditions are listed in **Supplementary Table 1.**

### Cell Lines

Human colon carcinoma Caco-2 cell line (ATCC#HTB37) was cultured in Dulbecco’s Modified Eagle’s Medium (DMEM; Thermo Fisher Scientific) containing 4 mM L-glutamine and 1 mM sodium pyruvate and supplemented with 10% Fetal Bovine Serum (FBS; Atlanta Biologicals). Caco-2 cells were seeded in a 12-well plate/trans-wells at 0.25 × 10^6^ cells/well. Cells were maintained in a CO_2_ incubator with 5% CO_2_ at 37°C and allowed to grow for 21-27 days ^78^ to form a well-differentiated monolayer with apical and basolateral surfaces. Caco-2 cell monolayers were used for adhesion, invasion, and translocation assays.

HEK-Dual™ hTLR2 (NF-κB-IL8) and HEK-Dual™ hTLR4 (NF-κB-SEAP) reporter cells (InVivoGen, San Diego, CA, USA) were grown in DMEM containing heat-inactivated 10% fetal bovine serum with antibiotics (Penicillin-Streptomycin, Normocin, Hygromycin B Gold and Zeocin) as per the vendor. Cells were incubated at 37°C in a humidified incubator with 5% CO_2_. A cell density of ∼5 × 10^5^ cells/mL was added to each well of a 96-well cell culture plate and incubated overnight at 37°C as above and treated with either bacteria or purified protein (see below).

### Sequence and Annotation of Lm H7014

A single colony of *Listeria monocytogenes* H7014 was cultured in TSBYE at 37°C for 16 h. DNA was isolated using Quick-DNA Fungal/Bacterial Microprep (Zymo, CA, USA). Qualified gDNA (A260/280 ∼1.8, A260/230 >2.0) was cleansed with RNAase before outsourcing for sequencing. Whole-genomic annotation was analyzed by Prokka (1.14.6) software and the Comprehensive Antibiotic Resistance Database (CARD) resistance gene identifier. The Lm F4244 sequence was used as a reference genome ^39^ for BLAST. The genomic maps and comparative genomics analysis were illustrated using Proksee (https://proksee.ca).

### Generation of *Listeria* Mutant Strains

The isogenic lap-deficient mutant (*lap*^─^) in strain H7014 was created using insertional mutagenesis as described ^25^. In brief, pPAad101 (a pAUL-A based temperature-sensitive shuttle vector) with partial *lap* (796 bp) was electroporated into Lm H7014 strain to replace the full *lap* at the exact location. Transformants were selected on Brain Heart Infusion agar (BHI; BD Bioscience) with erythromycin (Em; 5 µg/mL) at 30°C temperature for 48 h followed by subculture at a non-permissive temperature (42°C) overnight. The same procedure was used to create mInlA *lap*^─^. The H7014 *lap*-deficient complement strain (*lap*^─^ *lap*^+^) was constructed by introducing pMGS101 with a full *lap* (2601 bp). The positive clones were selected on BHI with erythromycin (Em; 5 µg/mL) and chloramphenicol (Cm; 5 µg/mL). The construction of H7014 *lap*^─^ and *lap*^─^ *lap*^+^ was confirmed through culturing them on modified Oxford agar (MOX) plates and Western blot for analysis of LAP, InlB, InlA, and LLO proteins.

### Extraction of Wall Teichoic Acid and PAGE

Wall teichoic acid (WTA) was extracted from each 40 mL bacterial culture as described before ^79^. Briefly, bacterial cells (OD600 adjusted to 1) were harvested and washed once with 30 ml of Buffer 1 [50 mM 2-(N-morpholino) ethanesulfonic acid (MES), pH 6.5] and then resuspended in 30 ml of Buffer 2 [4% (w/v) sodium dodecyl sulfate (SDS), 50 mM MES; pH 6.5]. Cells were boiled for 1 h, harvested by centrifugation (10,000 × g, 10 min), the pellets resuspended in Buffer 2, and then centrifuged (14,000 × g, 10 min). The pellet was washed once with Buffer 2, once with Buffer 3 [2% NaCl, 50 mM MES, pH 6.5], and finally with Buffer 1. Samples were treated with proteinase K buffer [20 mM Tris-HCl (pH 8.0), 0.5% (w/v) SDS, and 20 μg of proteinase K in 1 ml] and incubated at 50°C for ∼4 h. Samples were washed once with buffer 3, three times with distilled water to remove the SDS and resuspended in 1 mL of 0.1 M NaOH on a shaker at room temperature for 16 h to hydrolyze WTA. Insoluble cell wall debris was removed by centrifugation (14,000 × g, 10 min), and the supernatant containing the hydrolyzed WTA was either directly analyzed by non-denaturing PAGE (15%-acrylamide) followed by staining with Alcian blue and silver stain or stored at 4°C.

### Bacterial Protein Fractionation

Bacterial protein fractionation was done following the procedure described before ^25^. For preparation of the extracellular protein fraction, cell-free bacterial culture supernatants were precipitated with 60% ammonium sulfate at 4°C overnight on a roller. After centrifugation at 10,000×g for 15 min, precipitated protein pellets were resolubilized with 1 mL PBS containing 1× Halt Protease Inhibitor Cocktail (Thermo Scientific). The homogeneous solution was serially dialyzed against PBS using 12-kDa MWCO dialysis tubing to remove residual ammonia sulfate and kept at −20°C until use. For cell wall fraction (CWF), bacterial cell pellets were rinsed with PBS, and resuspended in cell wall extraction buffer (0.5% SDS, 20 mM Tris, pH 6.9) with 1× Halt Protease Inhibitor Cocktail, with rolling agitation at 37°C for 30 min. The samples were centrifuged at 16,000×g for 15 min at 4°C to obtain a clear supernatant and were designated the CWF. The samples were kept at −20°C until use. All cell wall fractions were tested for cytosolic contamination by aminopeptidase (PepC) assay ^33^ before use.

### Immunoblotting

All extracted fractions were pre-mixed with 4x Laemmli Sample Buffer [40% glycerol, 24% 1 M Tris HCl pH 6.8, 8% (w/v) SDS, 0.04% w/v bromophenol blue, 5% 2-mercaptoethanol (BME)] in 3:1 ratio and boiled for 10 min. The soluble fraction was loaded and separated on SDS-PAGE (10% acrylamide). Proteins were then transferred to 0.45 μm pore size Immobilon-P PVDF membranes (MilliporeSigma) using Trans-Blot electrophoresis apparatus (Bio-Rad Laboratories, Hercules, CA) at 35V for 12 h. Membranes were blocked with 5% non-fat dry milk (NFDM) or BSA prepared in Tris-buffered saline (TBS-T) (TBS; 20 mM Tris, 500 mM NaCl, 0.1% Tween-20 at pH 7.5; TBST) for one hour at room temperature. After washing with TBST, membranes were probed with the corresponding primary antibody (See **Supplementary Table 3**) in either 5% NFDM or BSA according to the manufacturer’s manual overnight at 4°C. After three more washes, membranes were probed with horseradish peroxidase-coupled (HRP) conjugated secondary antibody at half the concentration of the primary antibody. Protein bands were visualized by SignalFire ECL Reagent (CST, Danvers, MA). Images were acquired by ChemiDoc™ Imaging Systems (Bio-Rad Laboratories, Hercules, CA). Restore^TM^ Stripping Buffer was used to remove previously bound antibodies. Densitometry of protein bands was determined by ImageJ software (NIH) on at least three blots/biological replicates.

### RNA Isolation, Quality Control, Library Preparation, and Sequencing

Matured Caco-2 monolayer was infected with Lm F4244 WT or *lap*^─^ at MOI 10 following the adhesion procedure. Media supernatants were collected for a lactate dehydrogenase (LDH) assay to measure cell death. All samples had a cytotoxicity value lower than 5% (data not shown). After extensive washing, total RNA from Caco-2 was isolated using TRIzol reagent (Thermo Fisher Scientific) according to the manufacturer’s instructions. If RNA absorbance is greater than 2.0 at A260/280, the samples were treated with DNaseI to remove contaminated DNA. All clean total RNA was subject to Agilent 2100 bioanalyzer evaluation and determined to have an RNA integrity number (RIN) greater than 9. The following library preparation and sequencing was outsourced to BGI Groups (Shenzhen, China). Briefly, polyA tail mRNA/lncRNA was selected by oligo-dT-coated magnetic beads. The desired RNA was then fragmented before being converted into cDNA using reverse transcriptase with random hexamers. Only the sense-stranded RNA is amplified using dUTP method to avoid contamination of the second strand of cDNA. Stranded cDNA was ligated with an adapter before sequencing. The library was sequenced on DNBseq pair-end (PE) 100 bp with an outcome of at least 20 M clean reads.

### Sequence Bioinformatics Data Analysis

The differential gene expression (DGE) analysis was outsourced to SCGene (Guangzhou, China). All low-quality reads (adapter sequences, undetermined nucleotides, lower than Q20) were removed before reference annotation. StringTie was used to generate transcripts per million (TPM) ^80^. edgeR was used for DGE analysis.

### Translocation and Epithelial Permeability Assay

Caco-2 cells were seeded at 5 × 10^4^ cells/ml and grown as an intestinal epithelial-like monolayer for 14-21 days on Transwell inserts (3.0 μm pores, Corning). TEER was measured to monitor monolayer integrity (Millicells Voltmeter, Millipore). A TEER value of at least 400 Ω/cm^2^ was used as the basal value to monitor monolayer integrity. Stationary-phase bacterial cells were washed three times in PBS and resuspended in 500 μl DMEM (serum-free) and were applied on the apical side of the Transwell inserts at a multiplicity of infection (MOI) of 50. After 2 h for Lm and derivative strains at 37°C, 5% CO_2_, the media at the basolateral surface was aspirated, diluted, and plated on BHI-agar plates to enumerate translocated intracellular bacteria. For non-metabolizable 4-kDa, 40-kDa, and 70-kDa fluorescein isothiocyanate-dextran conjugate (FD4/40/70; 5 mg/mL, Sigma-Aldrich) flux analysis, FD was added to bacteria resuspended in DMEM and added to the apical side of the chamber. The basal chamber medium was collected, and fluorescence was measured (Em: 485 nm; Ex: 520 nm; Spectramax, Molecular Devices). To assay Lm-induced epithelial permeability, we monitored FD flux after 2, 8, and 16 h of incubation.

### Adhesion and Invasion Assay

Differentiated Caco-2 monolayers were washed with serum-free DMEM and infected with different strains of Lm and its derivative mutants at an MOI of 50 for 1 h and incubated in a 5% CO_2_ incubator at 37°C. For the adhesion assay, the monolayers were washed three times. For the invasion assay, cell monolayers were washed three times and then incubated with gentamycin (50 µg/mL) for 1 h to kill non-invaded Lm. Cells were washed thrice with sterile PBS to remove traces of loosely bound Lm and gentamycin. Cells from both adhesion and invasion experiments were treated with 0.1% Triton-X100 for 5-7 min, and the lysed cell suspensions were vortexed and serially diluted. Intracellular bacteria were enumerated by plating on BHI/TSAYE-agar plates.

### In Vitro TLR2 and TLR4 Signaling Assays

HEK-Dual™ hTLR2 cells (InVivogen) were cultured and maintained as described above in 96-well plates. For TLR2 inhibition assays, cells were pretreated with 100 µL of the TL2-C29 inhibitor (InvivoGen) at 200 µM at 37°C for 3 h in a humidified 5% CO₂ incubator. For TLR4 inhibition, CLI-95 was used at 1 µg/ml at 37°C for 3 h. Following pretreatment, 100 µL of purified proteins (LAP, InlA, and PGN @ 1 µg/ml; LPS @ 0.1µg/ml) or overnight-grown (∼18 h) bacterial cultures diluted to the desired concentration (to achieve MOI 10, 1) in DMEM was added to each well and incubated for an additional 6 or 24 h at 37°C. In addition, LAP and Hsp60 proteins were serially diluted and tested separately or in combination for dose-response EC_50_ ^81^ in the TLR2 reporter cell line. A 20 µL of HEK-Dual™ hTLR2 reporter cell culture supernatant was transferred to a new 96-well plate containing 180 µL of QUANTI-Blue™ solution (InvivoGen) to quantify secreted embryonic alkaline phosphatase (SEAP) activity, a measure of TLR2-dependent NF-κB/AP-1 activation. Plates were incubated for 1 h at 37°C in the dark, and absorbance was measured at 620 nm using an Epoch microplate spectrophotometer (BioTek Instruments).

### Protein Pull-Down Assay

Fifty microliters of Dynabeads™ (magnetic beads; Thermo-Fisher) were coated with 20-50 µg of each protein: LAP or InlA following the manufacturer’s instructions. The protein-coated beads were washed four times with 1x binding buffer (50 mM sodium phosphate, 300 mM NaCl, 0.01% Tween-20, pH 8) and mixed with HEK-Dual™ hTLR2 cell lysate in 1x Pull-down Buffer (3.25 mM sodium phosphate, 70 mM NaCl, 0.01% Tween-20, pH 7.4). Incubate the cell lysate-bead complexes at 4°C for 30 min in an end-to-end rotating shaker at 20 rpm. The beads were washed four times as before, resuspended in 25 μL 4× Laemmli sample buffer, and analyzed by SDS-PAGE (10% acrylamide) and Western blotting using anti-LAP mAb (mAb-H7), anti-Hsp60 pAb, and anti-InlA pAb (**Supplementary Table 3**).

### Mouse Experiment

All animal experiments were approved by the Purdue University Animal Care and Use Committee (IACUC Protocol no. 1201000595). Eight-to twelve-week-old (male or female) wild-type C57BL/6, *CD11B*^-/-^ (#003991, The Jackson Lab, ME, USA), *TLR2*^-/-^ (#004650, The Jackson Lab), *MLCK*^-/-^ (generously provided by Dr. Jerrold R. Turner, Harvard Medical School) from our breeding colony were used for the pathogenicity study. Mice receiving the same treatment were housed in the same cages, provided ad libitum feed and water, and acclimatized for at least 3 days before the experiments. Food was removed 4 - 6 h before oral gavage to prevent mechanical blockage of the *Listeria* inoculum by food in the stomach. *L. monocytogenes* and derivative mutant strains were suspended in 200 μL of PBS containing 1, 5 or 10 × 10^9^ CFU and were orally gavaged to randomly assigned mice using a stainless-steel ball-end feeding needle (Cadence Science). Naïve mice received the same amount of PBS. The food was returned immediately after the gavage. Mice were sacrificed 12-96 h post-gavage by CO_2_ asphyxiation.

### Establishing Gastroenteritis in a Mouse Model

To establish a mouse model of gastroenteritis, we designed time- and dose-dependent experiments using 8 to 12-week-old C57BL/6 mice. For the dose-dependent experiment, we orally gavaged mice with increasing doses (1×10^9^, 5×10^9^, or 1-2×10^10^ CFU/mouse) of wild-type strains F4244 and H7014 and assessed gastroenteritis symptoms, including fecal water content, gross morphological analysis of the colon, FD4 permeability, and inflammation. For the time-dependent experiment, Lm were orally gavaged @1-2×10^10^ CFU/mouse for 24, 48, 72 and 96 h. After each time point, mice were sacrificed and tested for gastroenteritis symptoms.

### Measurement of *In Vivo* Intestinal Permeability

Mice were orally gavaged with 4 or 40 kDa FITC dextran (15 mg/100 μL in PBS) 4 h before sacrifice. Blood was collected through cardiac puncture immediately after CO_2_ asphyxiation using a 21G needle and stored in serum-separating tubes (Thermo-Fisher). Fifty microliters of serum was mixed with an equal volume of PBS to measure FD levels. Urine excreted during CO_2_ asphyxiation was saved. Urine samples were diluted 1:10 in PBS (10 μL in 90 μL PBS) prior to FD measurement in a flat-bottom black 96 plate using SpectraMax (Ex: 490 nm and Em: 520 nm; Molecular Devices). The actual FD concentration was interpolated from an FD standard curve established for 1 to 2000 μg/mL.

### *Listeria monocytogenes* Burden Determination in Murine Organs and Tissues

The extra-intestinal tissues (liver, spleen, MLN) were harvested and homogenized by a tissue grinder in buffered *Listeria* enrichment broth (BLEB) immediately after euthanizing the mice ^31,82^. The intestinal sections (ileum, cecum, colon) were longitudinally incised to remove luminal contents and washed with PBS before suspending them in BLEB containing gentamicin (25 μg/ml) for 2 h at RT on a rotary shaker, washed 3 times, and homogenized to assess invaded Lm counts in intestinal tissues as described before ^31^. Samples were then serially diluted in PBS and plated on *Listeria* selective modified Oxford agar (MOX). A small section of the above organs was stored in 10% formalin for histopathology analysis.

### Gross Pathology and Histopathology

During the dissection of mice, images of intestinal organs and tissues including the cecum and colon were examined for fluid accumulation, fecal consistency, and gross pathology. Formalin-fixed intestinal and extra-intestinal tissue samples were stained with hematoxylin and eosin (H&E) and analyzed by a licensed veterinary pathologist. Inflammation was assessed based on histopathological morphological parameters, including the amount of polymorphonuclear leukocyte infiltration, mononuclear infiltration, and submucosal involvement. Each parameter was scored on a scale from 0 – 3. To determine the amount of polymorphonuclear and mononuclear leukocyte infiltration: 3, markedly increased; 2, moderately increased; 1, slightly increased; 0, normal. To grade the involvement of the submucosa: 3, 50% or greater of the submucosal diameter; 2, 10-50%; 1, less than 10%;0, normal. For the necrosis score of the liver and spleen: 3, massive necrosis; 2, more than 3 microscopic foci; 1, less than 3 microscopic foci. The inflammation scores of the liver and spleen: 2, moderate to marked inflammation associated with the foci of necrosis; 1, mild inflammation. The pathologist was blinded to sample identity.

### Fecal Water Content

Fecal water was measured using a previously described method ^83,84^ with some modifications. Briefly, feces from the distal 3 cm colon were squeezed out by forceps and collected in pre-weighed 1.5 mL microfuge tubes. The tubes with the feces were weighed and dried by incubating in a 55°C oven for 24 h. The tubes were reweighed, and the percentage decrease in fecal weight was determined.

### Diarrhea Scoring

The severity of diarrhea was assessed by the criteria outlined before ^83^. There are five grades of diarrhea, ranging from 1 to 5, where 1 is healthy and 5 is the most severe: 1, Normal colon with typically formed feces in the distal colon or rectum. 2, Cecum that is smaller than usual with areas of white and/or gas bubbles in the lumen; the rest of the colon is regular. 3, Severe typhlitis (white, shrunken cecum) with involvement of the proximal colon so that it is fluid-filled segments; formed stools in the distal colon. 4, Grade 3 plus fluid throughout most of the colon with soft feces in the distal colon. 5, No formed feces in the entire colon, and the cecum is pale and shrunken; the colon is filled with clear or bloody mucus.

### Cytokines and Lipocalin Assays

To determine proinflammatory mediator levels in the tissues, feces/intestinal tissue were resuspended in PBS (10 µL/mg) and vigorously vortexed with a bead beater (FastPrep-24 5G, MP Biomedicals) following the manufacturer’s instructions. The clear supernatants were used for quantification of proinflammatory cytokines/chemokines (RayBiotech, GA) and lipocalin-2 (R&D Systems, MN).

### RNA Preparation and qRT-PCR

Total RNA was isolated from the mouse ileal tissues using TRIzol reagent (Thermo Fisher Scientific) according to the manufacturer’s instructions. All RNA purity is greater than 2.0 at absorbance, A260/280. A 1 µg of total RNA was reverse transcribed to cDNA using NEB ProtoScript II Reverse Transcriptase (New England Biolabs, MA). Each transcript level was determined using Luna Universal qPCR Master Mix (New England Biolabs, MA) following manufacturer’s protocols. The thermal cycle condition is 95°C 10s, 50°C 30s, 72°C 30s, for 40 cycles. Primers were designed by Primer3Plus (https://www.primer3plus.com/) and synthesized by IDT (**Supplementary Table 3**). GAPDH was used as an internal control for each sample. The 2^-ΔΔCt^ method was used to calculate the relative changes in each gene expression.

### Isolation of Intestinal Cells and Flow Cytometry Analysis

Cells were isolated from both the small and large intestines. Peyer’s patches were removed from the small intestine before harvest. Intestines were cut longitudinally and washed with vigorous shaking 3 times with 40 mL of PBS to remove mucus. Washed intestinal tissues were then finely diced with scissors and transferred into digestion DMEM medium (Gibco, Thermo Fisher Scientific) containing 2% BSA, 10 μg/mL of DNase I (Roche), and 100 μg/mL of Liberase TL (Roche) or 1.5 mg/mL of type III collagenase (Worthington). Cells were incubated at 37°C for 1.5 h in a shaking incubator (250 rpm) before straining through a 100 μm nylon sieve and washing with PBS containing 2% fetal calf serum. The resulting cells were resuspended in 40% Percoll and centrifuged at 350g for 10 min at 4°C. Viable cells isolated from each tissue were enumerated via trypan blue staining and counted on a hemocytometer.

Antibodies used for flow cytometry are listed in **Supplementary Table 3**. Briefly, cells (1 - 2 × 10^6^) isolated from various tissues were preincubated with fixable viability dye (eBioscience, Thermo Fisher Scientific) and subsequently stained for surface markers (i.e., CD90.2, CD45). Cells were washed with FACS buffer (PBS, 0.1% BSA, 0.01% sodium azide) and fixed with 3.7% v/v formaldehyde in PBS for 20 minutes at 4°C.

### Transmission Electron Microscopy

Lm-infected Caco-2 monolayers were rinsed with PBS after designated incubation time and fixed with cold cacodylate buffer (2.5% Glutaraldehyde in 0.1 M cacodylate buffer, pH-7.4) at 4°C overnight. Samples were washed 4 times in the Cacodylate buffer, and incubated in Osmium solution (Osmium 1% + 0.8% FeCN) for 30 min. Samples were washed four times with DI water, incubated in 1% uranyl acetate for 30 min, rinsed three times in water, and sequentially dehydrated with 50, 75, 95% ethanol, 5 min each. Finally, samples were dehydrated four times (5 min each) with 100% ethanol, transitioned with acetonitrile twice, 5 min each time, followed by infiltration twice with 2:1 homogenized mixture of acetonitrile and resin, 15 min each time. Samples were incubated in 1:2 mixture of resin and acetonitrile for 10 min. Samples were incubated in Resin and DMP-30 mix for three times, 15 min each. Samples were kept in embedding frames and incubated overnight in the embedding oven. Sixty micrometer thin sections were cut and stained, followed by imaging on a Tecnai12 transmission electron microscope (TEM).

The TEM used for the mouse colon tissues followed a similar procedure with a brief modification. A 4 – 6 mm long portion of mouse colon was fixed in a cold cacodylate buffer (2.5% glutaraldehyde in 0.1 M cacodylate buffer) and kept at 4°C until use. The incubation with osmium tetroxide (Osmium tetroxide 1% + 0.8% FeCN) was increased to 2 h. The infiltration step with a homogenized mixture (2:1) of acetonitrile and resin was increased to overnight. The final incubation used a higher resin concentration (2% DMP-30 in resin) in the homogenized solution and was extended to 2 h.

### Confocal Immunofluorescence Microscopy

Ileum and colon tissues from different infection groups were fixed in 10% formalin for 48 h, and embedded in paraffin. Five µm-thick sections were prepared and deparaffinized using xylene, followed by an antigen retrieval process in sodium citrate buffer at sub-boiling temperature for 10 min. After cooling down, the samples were washed with PBS for 5 min and stained with the respective primary antibody overnight at 4°C. The samples were washed three times with PBS, 5 min each, followed by two two-hour incubations with fluorochrome-linked secondary antibodies at room temperature.

## Statistical Analysis

The experimental data were analyzed by GraphPad Prism version 11 (La Jolla, CA) using the Mann-Whitney test (two datasets) or one-way ANOVA with Tukey’s multiple comparisons (more than two datasets). All datasets have at least three independent/biological replicates, and the data are presented as the mean ± standard error of the mean (SEM) or as a box- and-whisker plot with the interquartile range.

TLR2 reporter responses were analyzed by ordinary two-way ANOVA to assess the effects of Hsp60 concentration, LAP concentration, and their interaction. Dunnett’s multiple-comparisons test was used to compare LAP-containing conditions with the corresponding LAP-free control at each Hsp60 concentration, with adjusted P values reported. Individual LAP and Hsp60 concentration-response curves were analyzed by nonlinear regression using a four-parameter logistic variable-slope model. Concentrations were converted to molar units using molecular masses of 94 kDa for LAP and 60 kDa for Hsp60. The Bottom was constrained to the experimentally determined untreated baseline, while Top, Hill slope, and EC_50_ (maximal-half concentration) were estimated by nonlinear regression. EC_50_ values with 95% confidence intervals were used to compare potency, and goodness of fit was assessed by R².

