## Supplemental Tables and Figures for "LAP-TLR2 Axis Drives Inflammatory Pathology during *Listeria monocytogenes* Gastroenteritis"

Dongqi Liu et al.

### Supplementary Tables

**Supplementary Table 1**. Gastroenteritis outbreak-causing *Listeria monocytogenes* strains used in this study.

| Strain | Year | Outbreak Associated | Location | Serotype | CC* | Source | Assembly |
|---|---|---|---|---|---|---|---|
| G4599 | 1993 | Yes | Italy | 1/2b | CC59 | Human | JPTX00000000 |
| G4600 | 1993 | Yes | Italy | 1/2b | CC59 | NA | PDT000158612.2 |
| G6006 | 1994 | Yes | USA | 1/2b | CC3 | Food (chocolate milk) | ASM2131255v1 |
| G6054 | 1994 | Yes | USA | 1/2b | CC3 | Human (stool) | ASM341783v1 |
| H7013 | 1997 | Yes | Italy | 4b | CC2 | Food (cold salad of corn & tuna) | PDT000158606.2 |
| H7014 | 1997 | Yes | Italy | 4b | CC2 | Human (stool & blood) | PDT000158626.2 |
| F4244 | 1991 | Yes | USA | 4b | CC6 | Human (CSF) | ASM274091v1 |

*Clonal Complex (CC) number is determined by Pasteur MLST ST. BIGSdb-Lm platform (http://bigsdb.pasteur.fr/listeria/)

**Supplementary Table 2**. *Listeria* genotypes and their growth conditions

| ***Listeria monocytogenes*** | **Genotype** | **Antibiotics used** | **Growth temperature** |
|---|---|---|---|
| F4244 | WT | - | 37°C |
| | *lap*$^{-}$ | Em5 | 42°C |
| | *lap*$^{-}$ *lap*$^{+}$ | Em5, Cm5 | 42°C |
| | Δ*inlA* | - | 37°C |
| | *mInlA* (murine InlA) | - | 37°C |
| | *mInlA lap*$^{-}$ | Em5 | 42°C |
| 10403S | Δ*hly* | - | 37°C |
| H7014 | WT | - | 37°C |
| | *lap*$^{-}$ | Em5 | 42°C |
| | *lap*$^{-}$ *lap*$^{+}$ | Em5, Cm5 | 42°C |
| G4600 | WT | - | 37°C |
| G6006 | WT | - | 37°C |
| G6054 | WT | - | 37°C |
| H7013 | WT | - | 37°C |

*Em5, 5 µg/mL of erythromycin; Cm5, 5 µg/mL of chloramphenicol.

**Supplementary Table 3.** Bacterial mutant strains, antibodies, cell lines, primers and reagents

| **REAGENT or RESOURCE** | **SOURCE** | **IDENTIFIER** |
|---|---|---|
| **Bacterial Strains** | | |
| *L. monocytogenes* F4244 serovar 4b (WT), CC6 | Our collection [1] | N/A |
| *L. monocytogenes* F4244 *lap*⁻ (isogenic *lap* insertion mutant) | Kim et al., 2006 [2] | N/A |
| *L. monocytogenes* F4244 *lap*⁻ *lap*⁺ (*lap* complemented) | Kim et al., 2006 [2] | N/A |
| *L. monocytogenes* F4244 Δ*inlA* (in-frame deletion) | Burkholder and Bhunia, 2010 [3] | N/A |
| *L. monocytogenes* F4244 *mInlA* (murinized InlA strain) | Bai et al. 2021 [4] | N/A |
| *L. monocytogenes* F4244 *mInlA lap*⁻ (isogenic *lap* insertion mutant in murinized InlA background) | Drolia et al. 2024 [5] | N/A |
| *Listeria monocytogenes* 10403S Δ*hly* (in-frame deletion) | Glomski et al. 2003 [6] | DP-L2161 |
| *L. monocytogenes* H7014 serovar 4b (WT), CC2, Italy, 1997 | CDC | N/A |
| *L. monocytogenes* H7014 *lap*⁻ (isogenic *lap* insertion mutant) | This study | N/A |
| *L. monocytogenes* H7014 *lap*⁻ *lap*⁺ (*lap* complemented) | This study | N/A |
| *L. monocytogenes* H7013 serovar 4b, Italy, 1997 | CDC | N/A |
| *L. monocytogenes* G4600 serovar 1/2b, Italy, 1993 | CDC | N/A |
| *L. monocytogenes* G6006 serovar 1/2b, USA, 1994 | CDC | N/A |
| *L. monocytogenes* G6054 serovar 1/2b, USA, 1994 | CDC | N/A |
| *E. coli* DH5α pMSG101-LAP | [7] | N/A |
| *E. coli* DH5α pAUL-AAD | [7] | N/A |
| **Mammalian Cell Lines** | | |
| Caco-2 | ATCC | Cat# HTB37 |
| HEK-Blue-Lucia hTLR2 cell | Invivogen | hkd-htlr2ni |
| **Antibodies** | | |
| Mouse monoclonal α-LAP (EM-H7) | Our Lab [8] | NA |
| Rabbit polyclonal α-InlA | Our Lab [3] | NA |
| Rabbit polyclonal α-InlB (Lm404) | Our Lab [9,10] | NA |
| Rabbit polyclonal α-LLO | This study | NA |
| Goat anti-rabbit HRP-conjugated IgG | Cell Signaling Technology | Cat # 7074, RRID: AB_2099233 |
| Horse anti-mouse HRP-conjugated IgG | Cell Signaling Technology | Cat # 7076, RRID: AB_330924 |
| α-CD90.2 (Thy1.2)-Pacific Blue | Biolegend | Cat#105323 |
| α-CD45-BV711 | Biolegend | Cat#103147 |
| α-Ly6G-AlexaFluor488 | Biolegend | Cat#127625 |
| α-EpCAM (CD326)-PerCP Cyanine5.5 | Biolegend | Cat#118219 |
| α-B220 (CD45R)-PE | Biolegend | Cat#103207 |
| α-CCR2-PE Cyanine 5 | Biolegend | Cat#150637 |
| α-Ly6C-PE Cyanine 7 | Biolegend | Cat#128017 |
| α-CD11c-APC | Biolegend | Cat#117309 |
| eFluor 780 (Viability Dye) | eBioscience | Cat#65-0865-14 |

| Commercial Reagents and Chemicals | | |
|---|---|---|
| Pierce™ BCA Protein Assay Kit | Thermo Fisher | Cat# 23225 |
| Halt™ Protease Inhibitor Cocktail (100X) | Thermo Fisher | Cat# 78429 |
| PMSF Protease Inhibitor | Thermo Fisher | Cat# 36978 |
| Halt™ Protease and Phosphatase Inhibitor Cocktail (100X) | Thermo Fisher | Cat# 78440 |
| M-PER™ Mammalian Protein Extraction Reagent | Thermo Fisher | Cat# 78501 |
| Phusion® High-Fidelity DNA Polymerase | NEB | Cat# M0530 |
| Mouse Lipocalin-2/NGAL Duoset ELISA | R&D | Cat# DY1857 |
| Fluorescent isothiocyanate-dextran 4 kDa | MillporeSigma | Cat# FD4 |
| Fluorescent isothiocyanate-dextran 40 kDa | MillporeSigma | Cat# FD40 |
| Fluorescent isothiocyanate-dextran 70 kDa | MillporeSigma | Cat# FD70 |
| RNAlater | Thermo Fisher | Cat# AM7021 |
| TL2-C29 (TLR2 Inhibitor) | Invivogen | Inh-c29 |
| Modified Oxford agar Base | Neogen Corporation | Cat# 7428 |
| Modified Oxford agar Base supplement | Neogen Corporation | Cat# 7991 |
| Buffered Listeria enrichment broth | Neogen | Cat# 7675 |
| Buffered Listeria enrichment broth supplement | Neogen | Cat# 7980 |
| BD Microgard™ Gold | BD | Cat# 365967 |
| **Oligonucleotides** | | |
| LAP_NcoI_F CGCCCATGGCAATTAAAGAAAATGC | This study | N/A |
| LAP_XhoI_R CGCCTCGAGTTAAACACCTTTGTAAG | This study | N/A |
| mNHE2_F GGACACTGGGGTCACAACTT | This study | N/A |
| mNHE2_R GCGGTTTGAGCTGTTGTCTT | This study | N/A |
| mNHE3_F CCTTCATTCGCTCCCCAAGT | This study | N/A |
| mNHE3_R GTGTTAGCTCGTGCCGACTA | This study | N/A |
| mSGLT1_F TTCCAGACGTGCACCTGTAC | This study | N/A |
| mSGLT1_R GCACACTGGCTTCAAATGCA | This study | N/A |
| mDRA_F ACGAACATCGCAGACCTTGT | This study | N/A |
| mDRA_R TGTAGCGGCTGCAGAAGAAA | This study | N/A |
| mCLCA1_F GAATGAACCCACCACGTCCT | This study | N/A |
| mCLCA1_R TTGAGTCTTCGGGAATGGGC | This study | N/A |
| mCFTR_F AAAGTTGTCAAGCCGCGTTC | This study | N/A |
| mCFTR_R TATCTGTACGGCAGTGGGGA | This study | N/A |
| mNKCC1_F GGCTGCGTTGCTCACATATG | This study | N/A |
| mNKCC1_R GGCGGATAACCACAACTCCA | This study | N/A |
| mGAPDH_F TTCACCACCATGGAGAAGGC | This study | N/A |
| mGAPDH_R TGAAGTCGCAGGAGACAACC | This study | N/A |
| **Analysis Software** | | |
| ImageJ | NIH | Ver 1.53k |
| GraphPad Prism | Graph Pad | Version 8.4.3 |
| SnapGene | GSL Biotech | Version 6.0.2 |
| Proksee | Genome Canada/Genome Alberta | NA |

#### Supplementary Figures

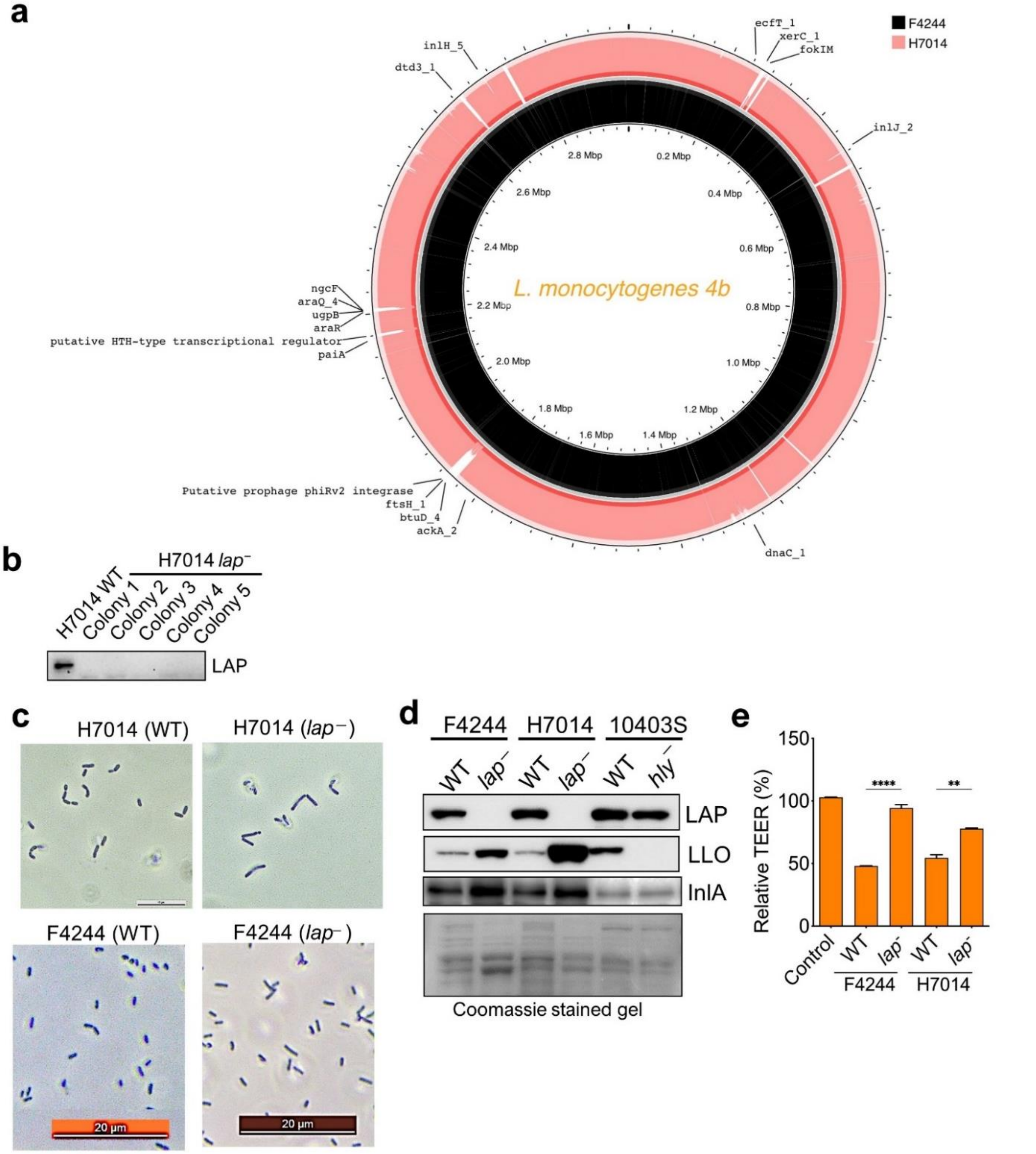


**Supplementary Figure 1. LAP Mediates Epithelial Barrier Disruption and Lm Translocation by Expanding Paracellular Gap**

**a.** Circular genomic alignment and comparative genomics between Lm F4244 (Black) and H7014 (Pink). **b,c.** Immunoblot (b) and microscopic morphology (c) characterization of the isogenic *lap*-deficient strain generated from the H7014 background. **d.** Immunoblots of LAP, InlA, and LLO levels in Lm WT, *lap*⁻ strains (F4244 and H7014 backgrounds). Lm 10403S WT and *hly*⁻ (hemolysin-deficient) strains serve as positive and negative controls for LLO expression, respectively. Coomassie blue staining of total protein was used as a loading control. **e.** Relative TEER values after exposure to bacteria in a Transwell setup.

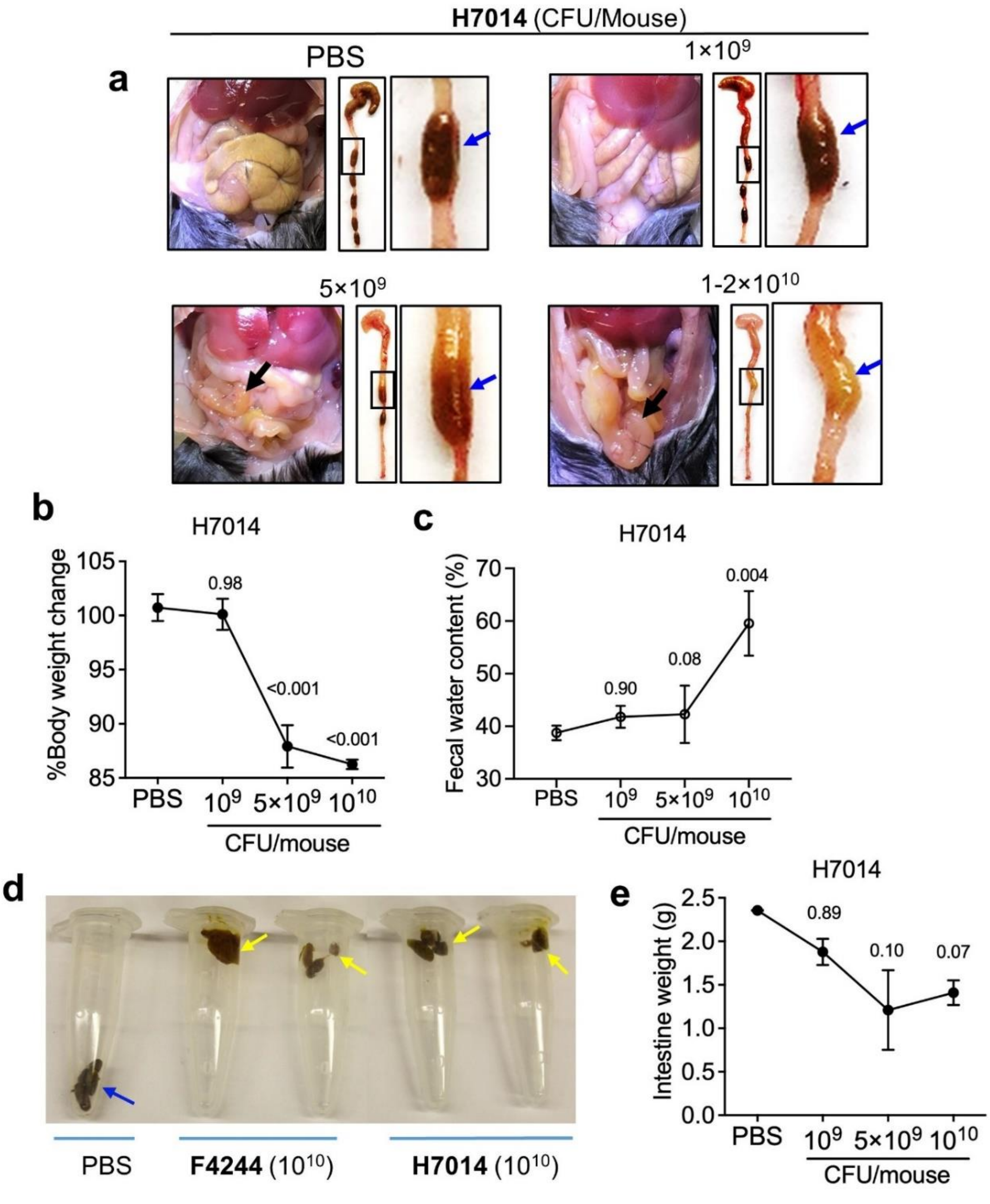


**Supplementary Figure 2. Acute Gastroenteritis in mice gavaged with Lm.**

**a.** Representative images of gross intestinal pathology of C57BL/6 WT mice that were intragastrically infected with 1 x $10^9$, 5 x $10^9$, 1-2 x $10^{10}$ CFU/mouse of Lm H7014, or PBS (Mock). **b.** Percentage change in total body weight. **c.** Analysis of fecal water content. PBS: n=6, Lm-infected mice: n=12-15

**d.** Fecal consistency (stickiness) after dosing with 1-2 x $10^{10}$ CFU/mouse of F4244 and H7014 after 48 hpi. **e.** Intestinal weight of mice infected with Lm H7014 after 48 h. n=5-6. All error bars represent SEM, marked with P values. All experiments were repeated at least three times unless otherwise noted.

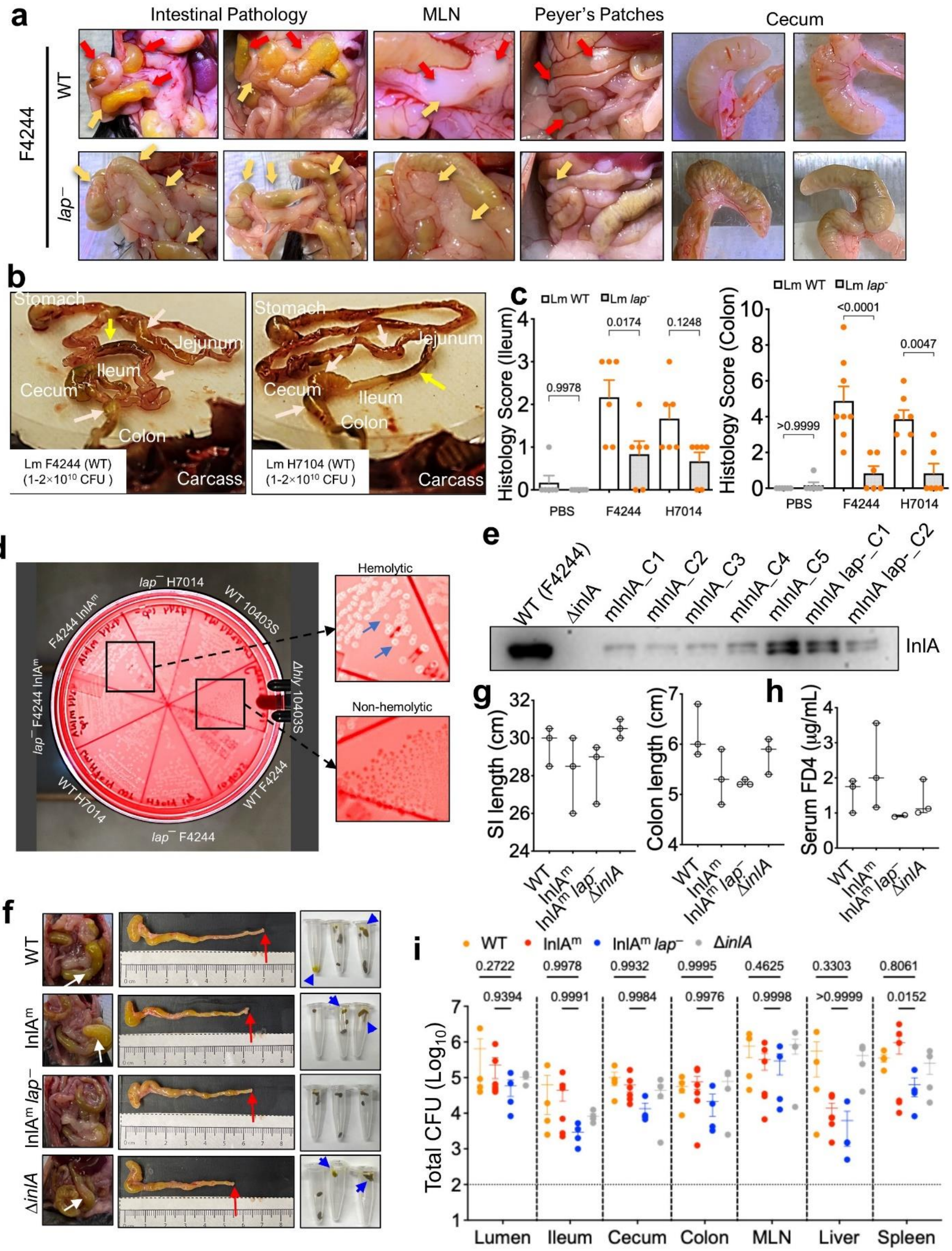

**Supplementary Figure 3. LAP is the Main Driver of Gastroenteritis In Vivo.**
**a.** Representative images of gross intestinal pathology and close-up images of mesenteric lymph nodes (MLN), Peyer's patches, and cecal contents from mice infected with Lm F4244 WT or *lap*⁻.
**b.** Representative images of the whole intestine from stomach to colon infected with Lm F4244 WT or *lap*⁻.
**c.** Intestinal histopathological scores of Lm WT or *lap*⁻ in both F4244 and H7014 backgrounds.
**d.** Characterization of Lm isolates and variants on blood agar. All tested strains, with the exception of the10403S Δ*hly*, exhibited clear zones of $\boldsymbol{\beta}$-hemolysis (blue arrows from zoom-in image).
**e.** Immunoblot confirming InlA expression in F4244 $InlA^m$ and $InlA^m$ *lap*⁻ strains. F4244 WT and ΔinlA serve as positive and negative controls, respectively.
**f-i**. Assessment of gross intestinal pathology and fecal morphology (f), small and colon length of mice infected with Lm F4244 WT, $InlA^m$, $InlA^m$ *lap*⁻, ΔInlA (g), serum FD4 permeability (h), and tissue Lm burdens (i) in mice infected with Lm InlA variants. The symptoms of $InlA^m$ *lap*⁻ are less intense, characterized by visible pellet structure (chunks) rather than watery stool; reduced mucus residues on the pellets compared to WT and $InlA^m$ (blue arrows). B6 mice were intragastrically gavaged with 1-2 x $10^{10}$ CFU/mice Lm F4244 WT, murinized InlA ($InlA^m$), $InlA^m$ *lap*⁻, and ΔInlA.

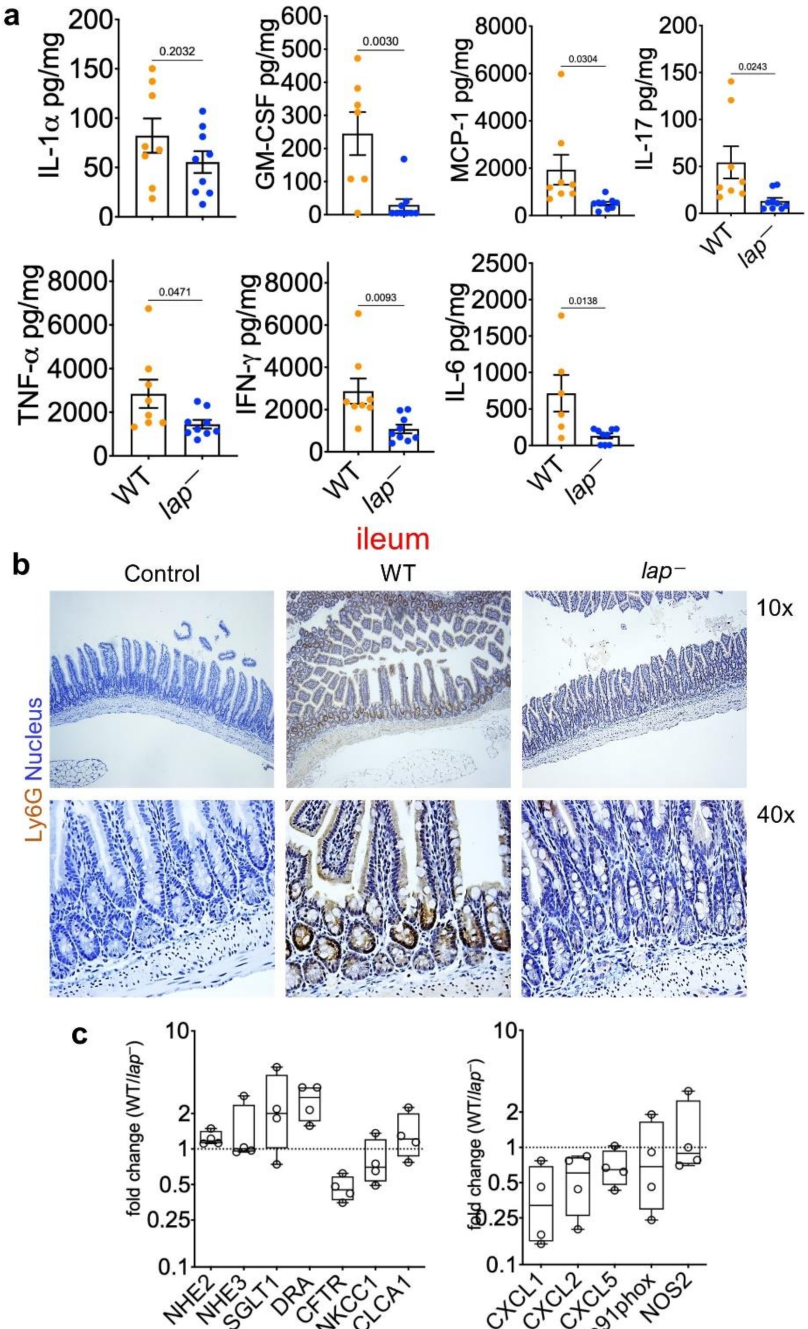


**Supplementary Figure 4. LAP-induced leaky gut exacerbates gastroenteritis symptoms through increased epithelial cell death and immune cell infiltration**

**a.** Colonic proinflammatory cytokine profiling of mice infected with Lm H7014 WT and Lm H7014 *lap*⁻. **b.** Representative IHC staining for Ly6G (brown) in the ileum. Lm background is F4244. **c**. Relative mRNA levels of ion absorptive/secretory ion channels for Lm WT/ *lap*⁻.

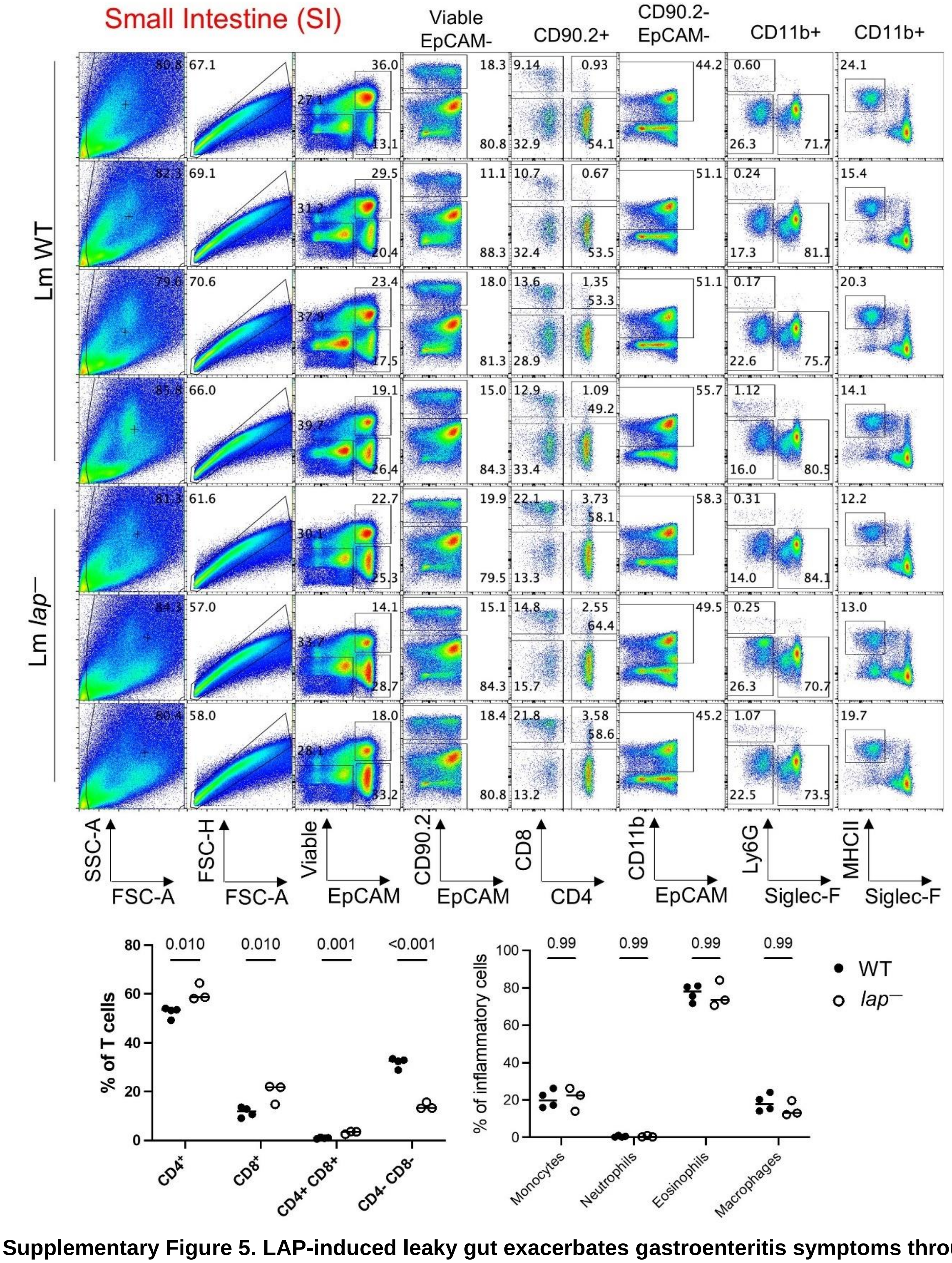


**Supplementary Figure 5. LAP-induced leaky gut exacerbates gastroenteritis symptoms through increased epithelial cell death and immune cell infiltration**. Flow cytometry gating strategy and immune cell subset frequencies in the small intestine. B6 mice were infected with Lm F4244 WT or Lm F4244 *lap*⁻ at 1-2 x$10^{10}$ CFU/mouse.

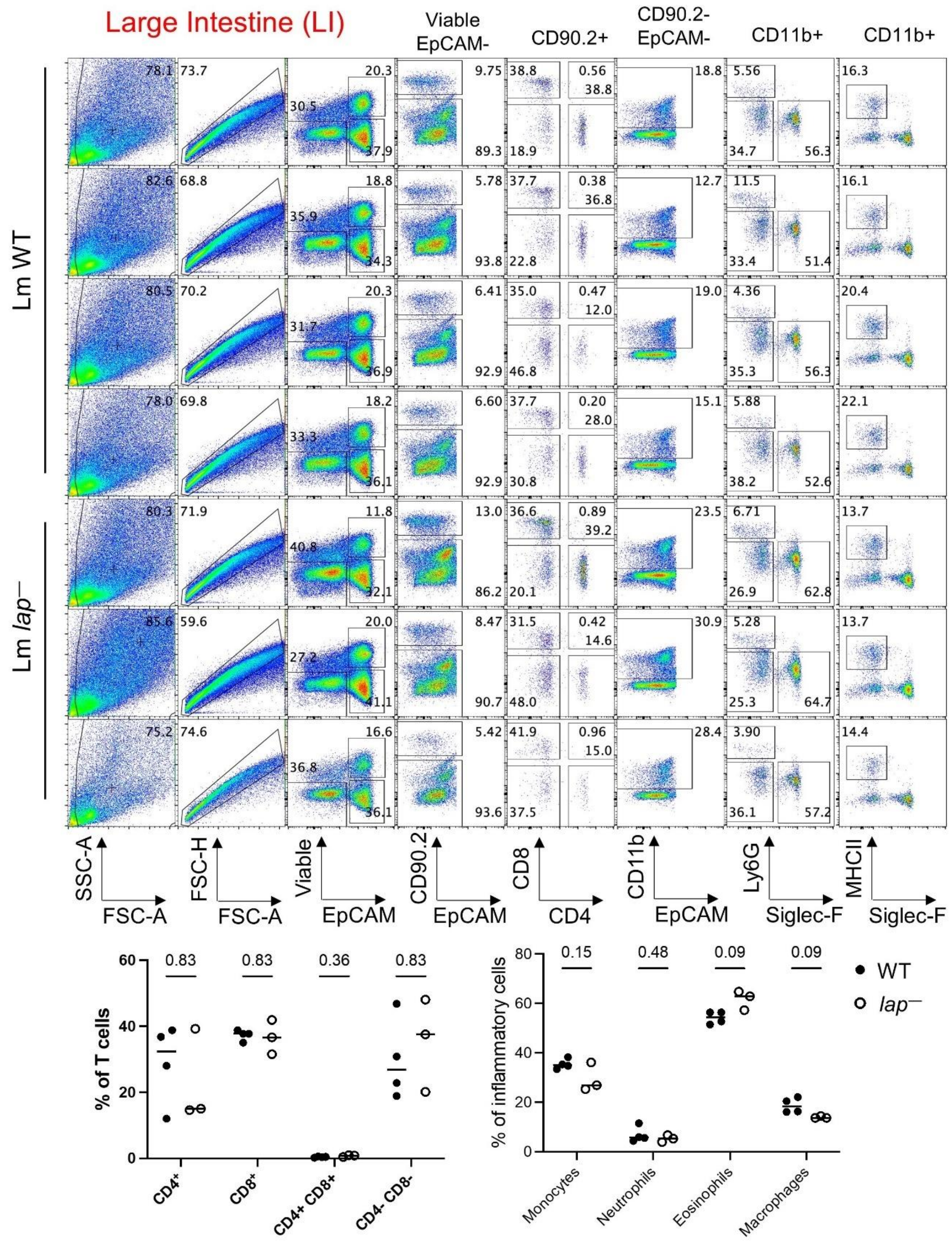


**Supplementary Figure 6. LAP-induced leaky gut exacerbates gastroenteritis symptoms through increased epithelial cell death and immune cell infiltration.** Flow cytometry gating strategy and immune cell subset frequencies in the large intestine. B6 mice were infected with Lm F4244 WT or Lm F4244 *lap*⁻ at 1-2 x$10^{10}$ CFU/mouse.

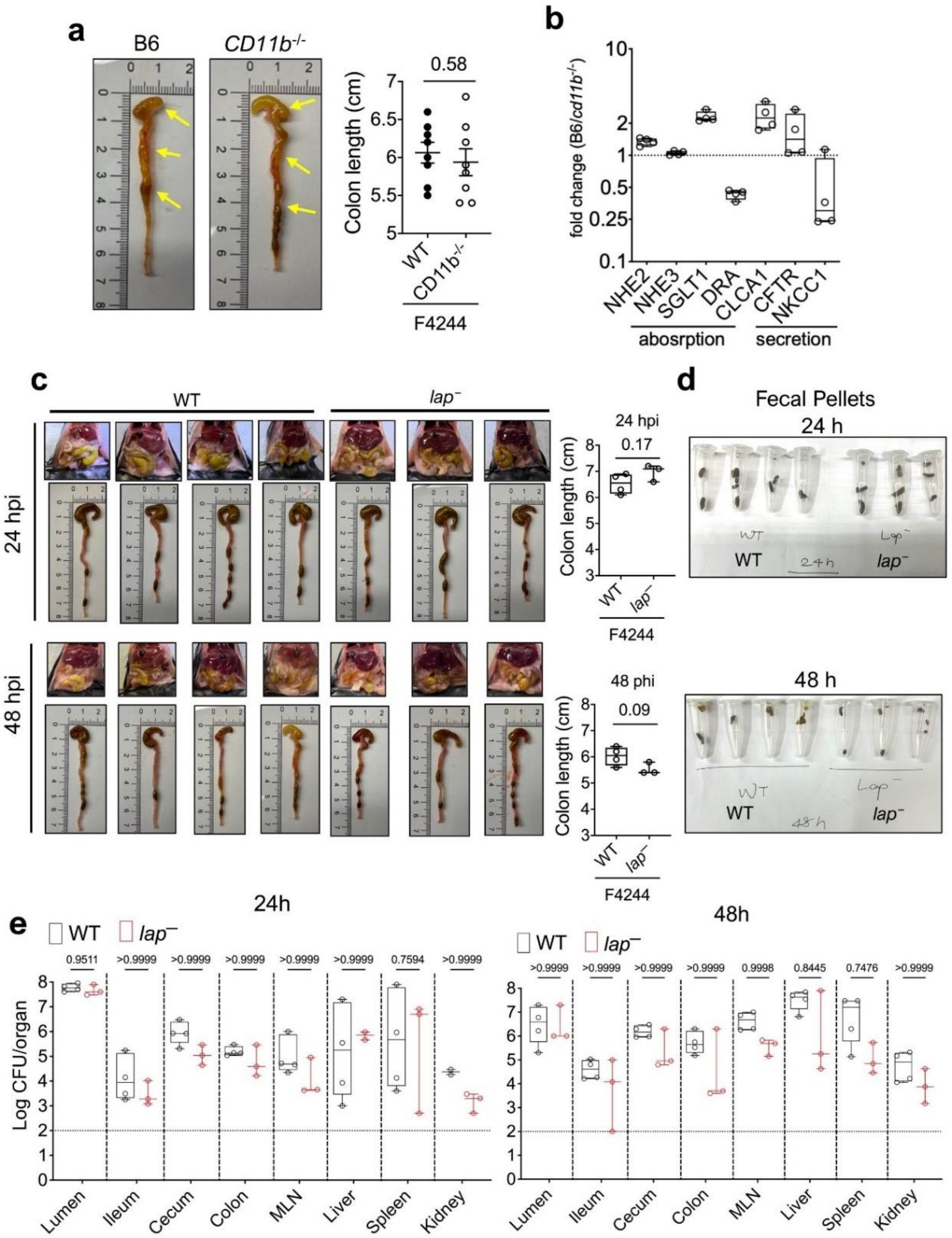


**Supplementary Fig 7. CD11b+ Myeloid Cells Mediate Diarrheal Symptoms and Contain Lm Spread in the intestine.**

**a.** Representative images and corresponding bar plots showing the length of the small (a)(stomach-ileum) and large intestine (b)(cecum-colon). Percentage of viable colonic IEC cells from Lm-infected B6 and Cd11b$^{-/-}$ mice. **b.** Relative mRNA level of ion absorptive/secretory ion channels and $Ca^{2+}$ dependent ion channel regulator from the ileum. **c.** Gross intestinal pathology and cecum-colon length of *CD11b*$^{-/-}$ mice infected with 1 x $10^{10}$ CFU/mouse F4244 Lm WT or *lap*$^{-}$ at 24 and 48 hpi. Colon length after 24 and 48 h. **d.** Fecal morphology and volume after 24 and 48 h. **e.** Lm burden in the intestinal and extra-intestinal tissues.

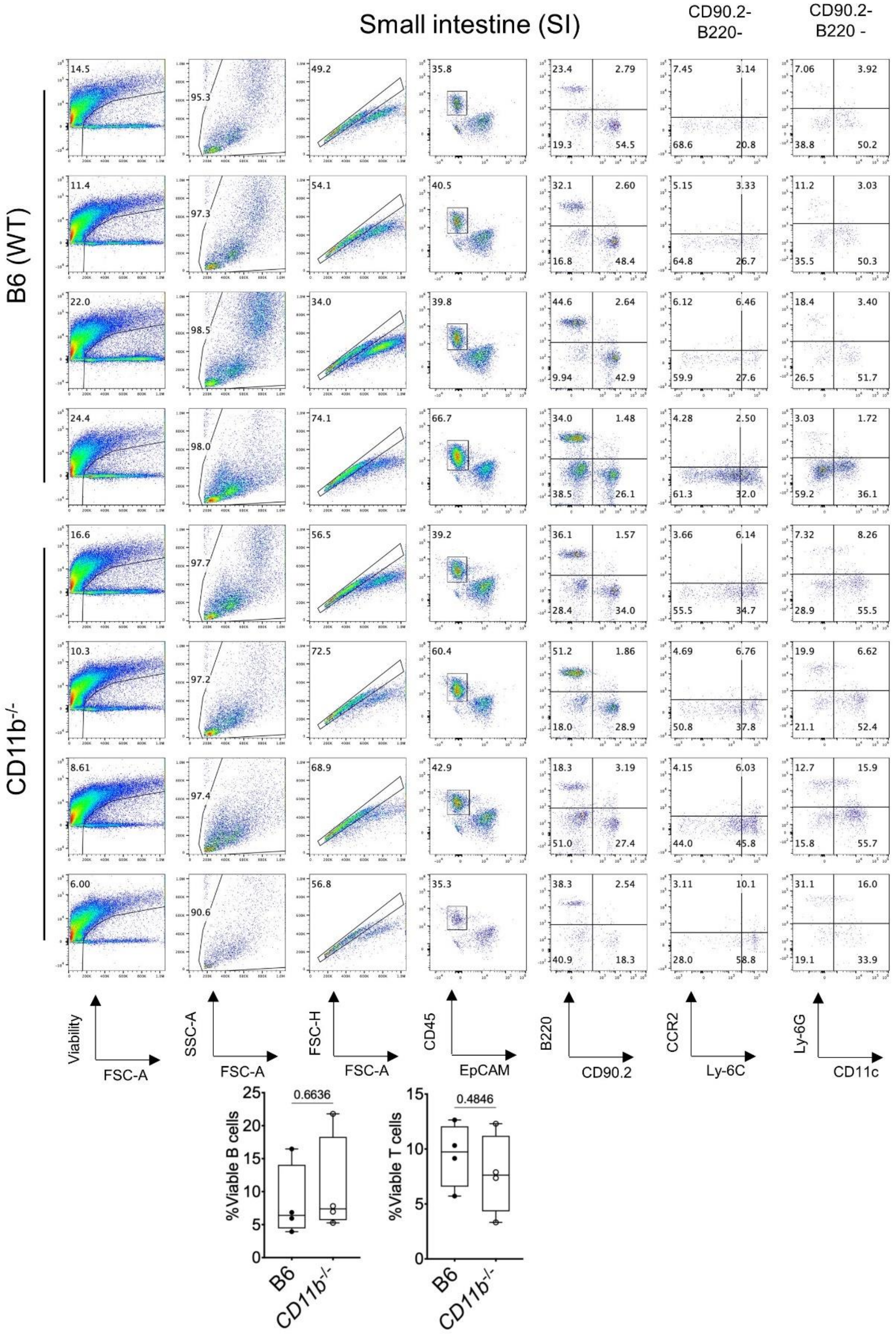


**Supplementary Fig 8. Flow cytometry gating strategy and immune cell subset frequencies** in the small intestine of *CD11b*$^{-/-}$ mice. Animals were infected with Lm F4244 WT at 1-2 x$10^{10}$ CFU/mouse.

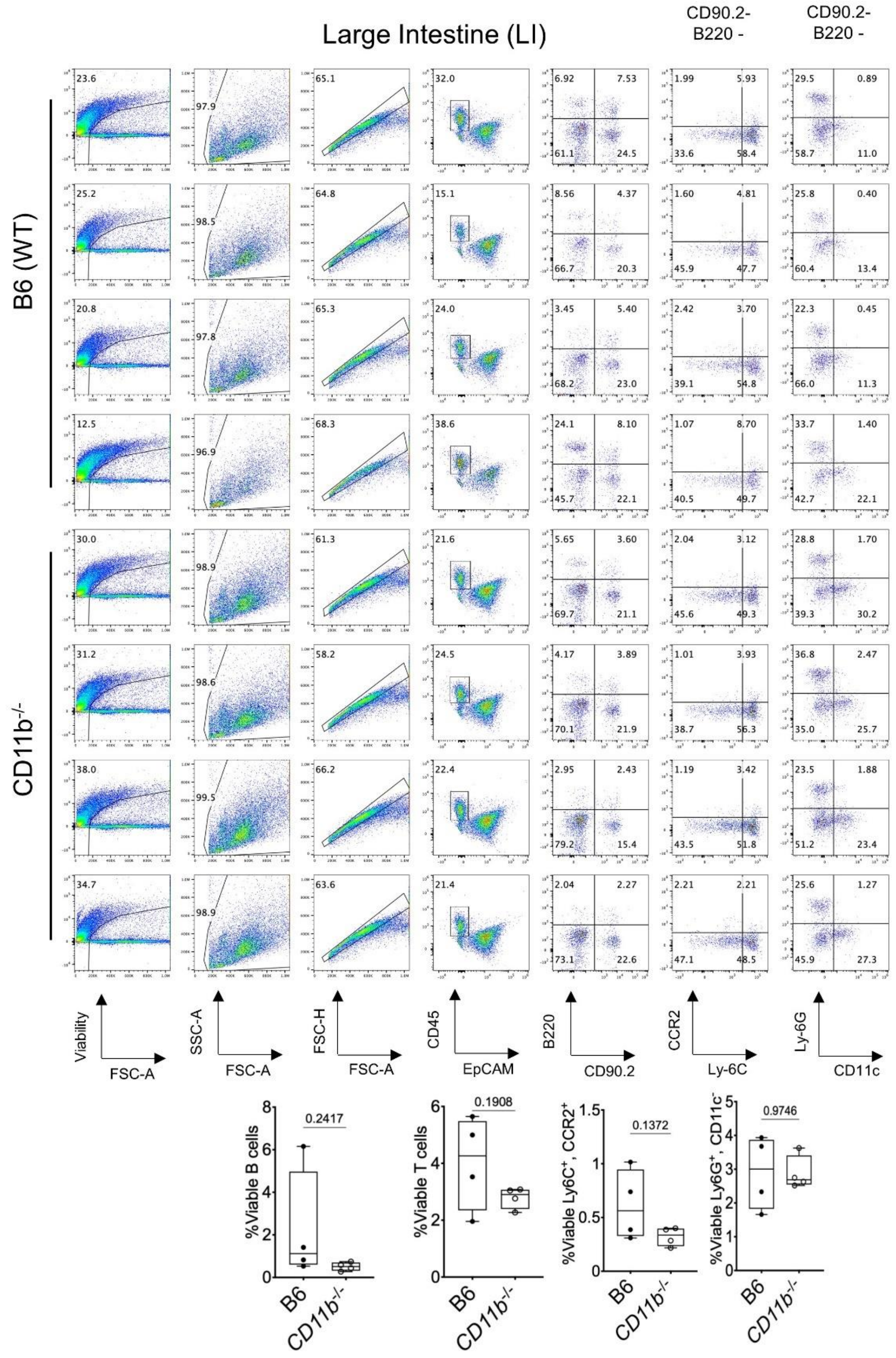


**Supplementary Fig 9. Flow cytometry gating strategy and immune cell subset frequencies** in the large intestine of *CD11b*$^{-/-}$ mice. Animals were infected with Lm F4244 WT at 1-2 x$10^{10}$ CFU/mouse.

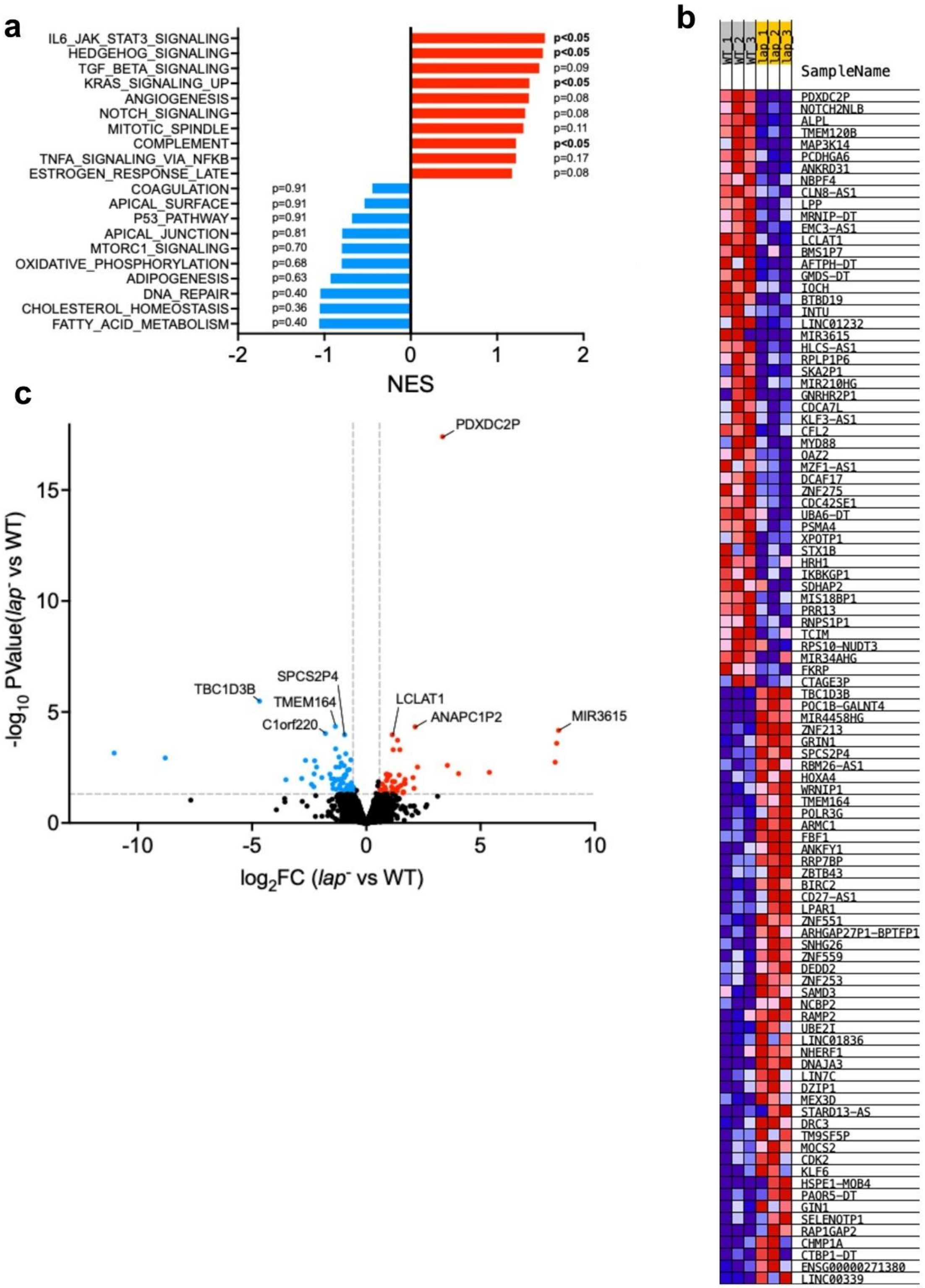


**Supplementary Fig. 10. LAP-Induced Gastroenteritis is Mediated by TLR2 Signaling.**
**a.** GESA analysis of top 10 upregulated and downregulated pathways. Significantly different pathways were bolded. **b.** RNAseq heatmap illustrating the complete list of differentially expressed genes (DEGs) in Caco-2 cells infected with Lm WT compared to the *lap*⁻ strain. **c.** Volcano plot of DEGs.

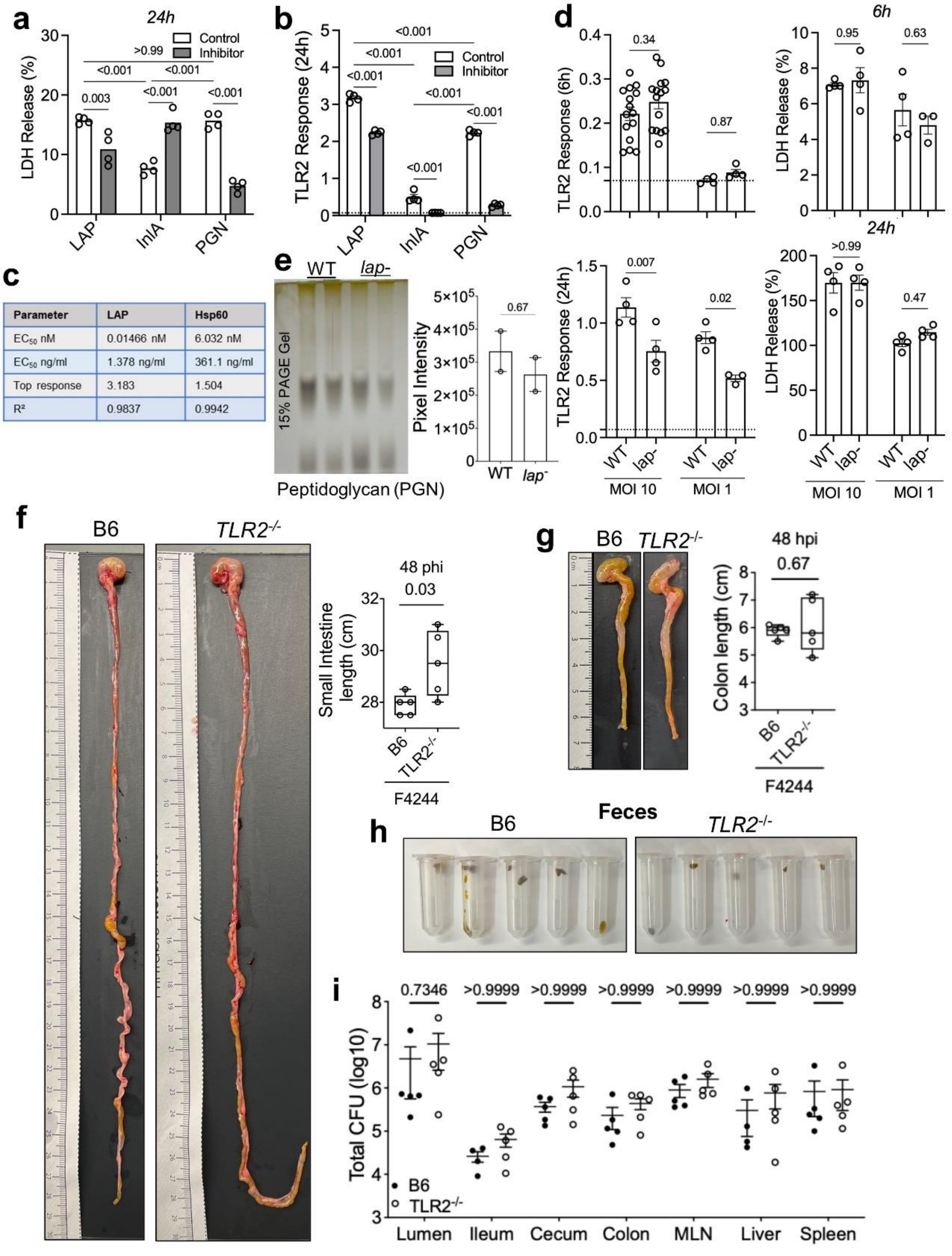


**Supplementary Figure 11. LAP-Induced Gastroenteritis is Mediated by TLR2 Signaling.**

**a.** LDH cell toxicity assay for hHEK-TLR2 reporter cell line treated with purified proteins.

**b.** TLR2 signal responses in hHEK-TLR2 reporter cell line with LAP, InlA and PGN and TLR2 inhibitor (TL2-C29) after 24 h.

**c**. TLR2 signal response ($EC_{50}$ estimation) to variable doses of LAP and Hsp60.
**d.** TLR2 signal responses in hHEK-TLR2 reporter cell line treated with live Lm WT or *lap*⁻ at 6 h and 24 h post-exposure and corresponding LDH release assay.
**e.** Wall teichoic acid (WTA) profiles of Lm WT and *lap*⁻ analyzed by non-denaturing PAGE with Alcian blue staining (top) and corresponding densitometric pixel intensity analysis (bottom).
**f,g.** Representative images and corresponding bar plots quantifying the length of the small (stomach-ileum) (e) and large intestine (cecum-colon) (f).
**h.** Representative images of fecal pellets of mice infected with Lm collected at 48 hpi.
**i.** Lm burden in intestinal and extra-intestinal tissues at 48 hpi.

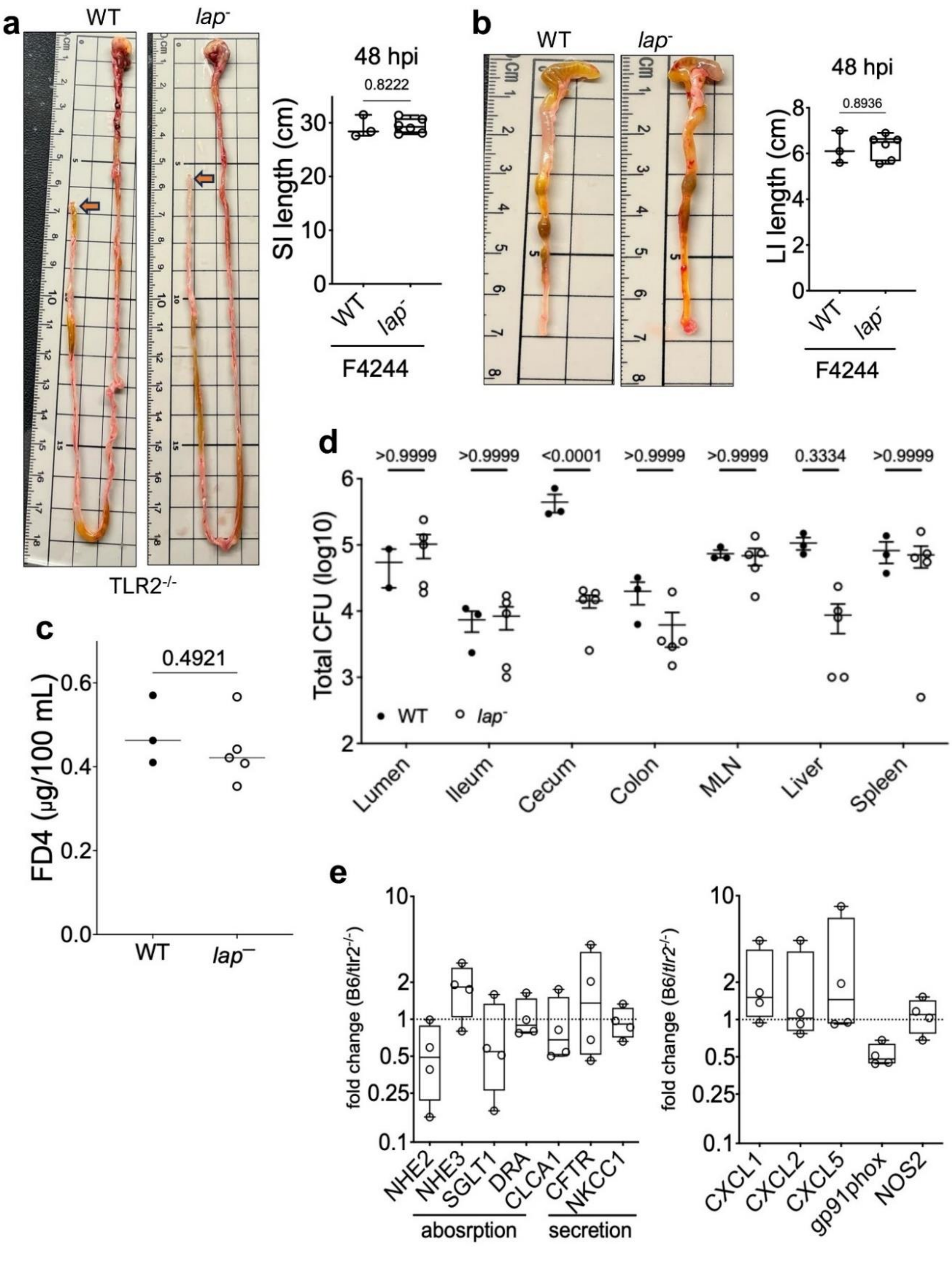


**Supplementary Figure 12. LAP-Induced Gastroenteritis is Mediated by TLR2 Signaling.**
**a,b.** Representative images and corresponding bar plots quantifying the length of the small (stomach-ileum) intestine (a) and large intestine (cecum-colon) (b) from *TLR2*$^{-/-}$ mice infected with Lm WT and the *lap*$^{-}$ strain. **c.** FD4 permeability in *TLR2*$^{-/-}$ mice infected with Lm WT and the *lap*$^{-}$ strain. **d.** Lm burden in intestinal and extra-intestinal tissue of *TLR2*$^{-/-}$ mice at 48 phi. **e.** Relative mRNA expression levels of ion channels and inflammatory markers from the ileum.

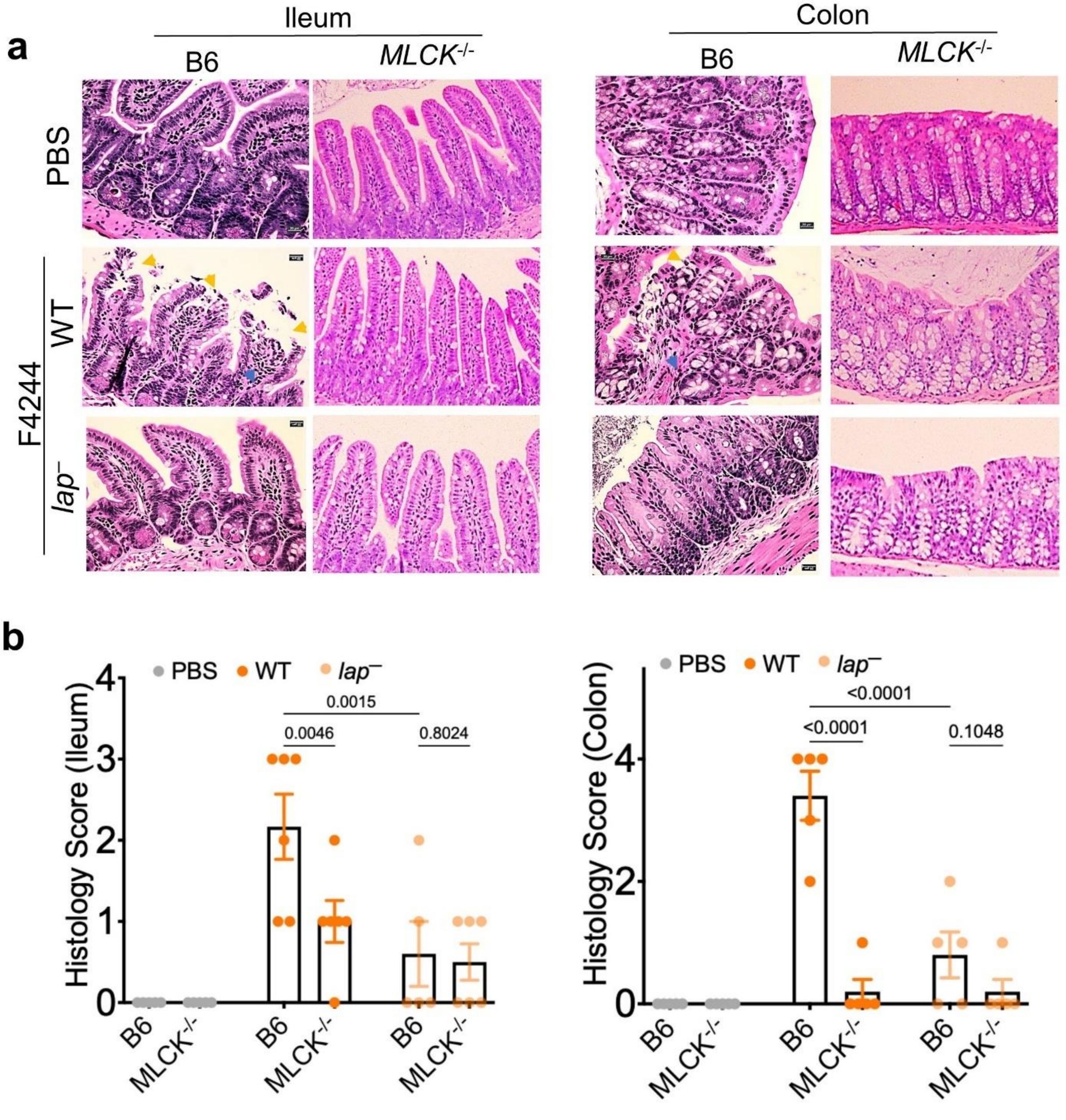


**Supplementary Figure 13. MLCK deficiency preserves the paracellular gap and attenuates LAP-induced gastroenteritis.**

**a.b.** Representative histopathology images (a) and blinded scoring data (b) from ileal and colonic tissues from Lm WT and *lap*$^{-}$ infected B6 and MLCK$^{-/-}$ mice at 48 hpi. All error bars represent SEM and are marked with p-values. All experiments were repeated at least three times unless otherwise noted.

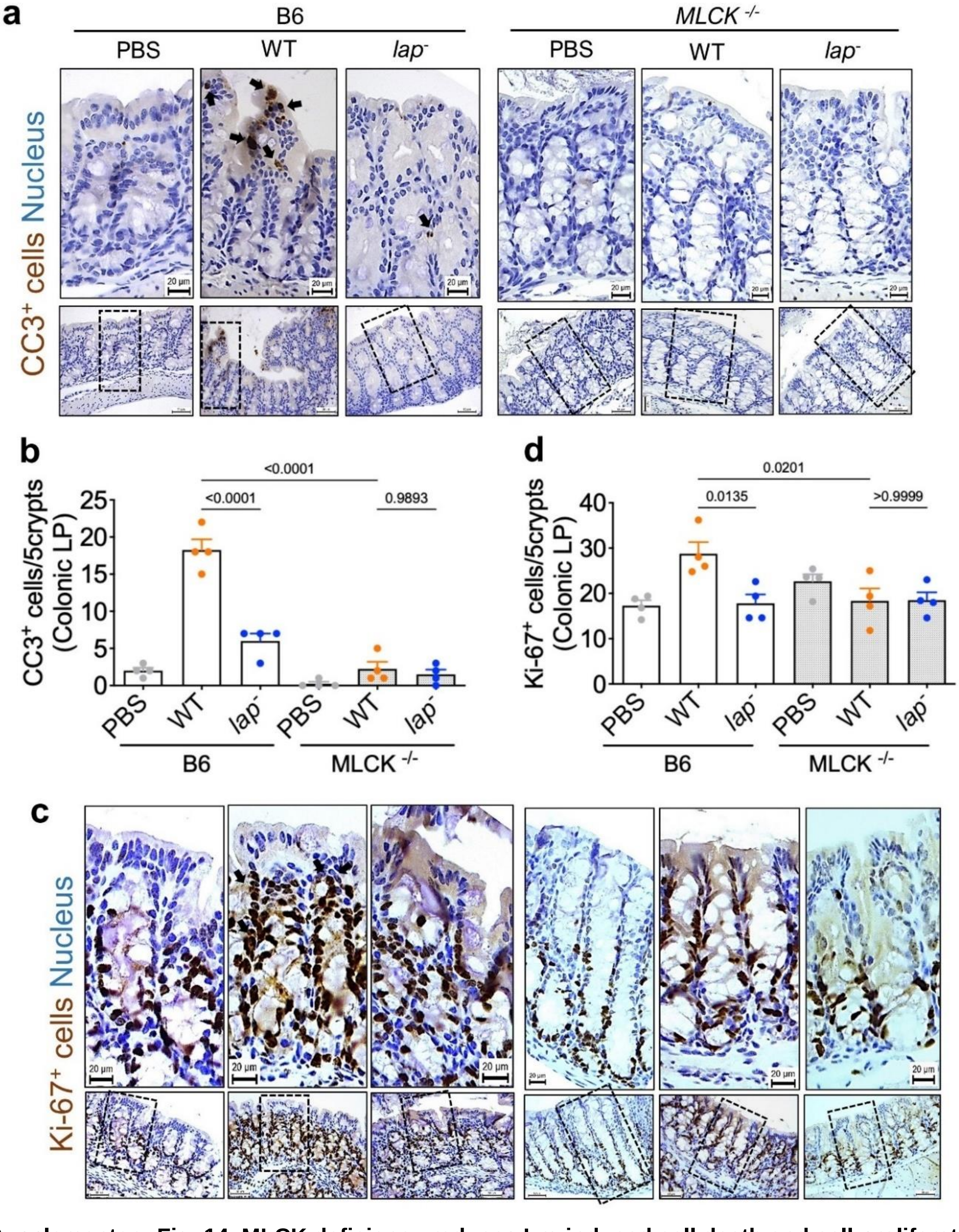


**Supplementary Fig. 14. MLCK deficiency reduces Lm-induced cell death and cell proliferation at the lamina propria.**

**a.b.** Representative colonic IHC staining of CC3⁺ (Cleaved caspase 3) at the lamina propria and quantitative analysis showing the mean counts of CC3⁺ (b). The nucleus was counterstained with hematoxylin. **c.d.** Representative colonic IHC staining of Ki67⁺ at the lamina propria and quantitative analysis showing the mean counts of Ki67⁺ (d) cells within the lamina propria. The nucleus was counterstained with hematoxylin. All error bars represent SEM and are marked with p values. All experiments were repeated at least three times unless otherwise noted.

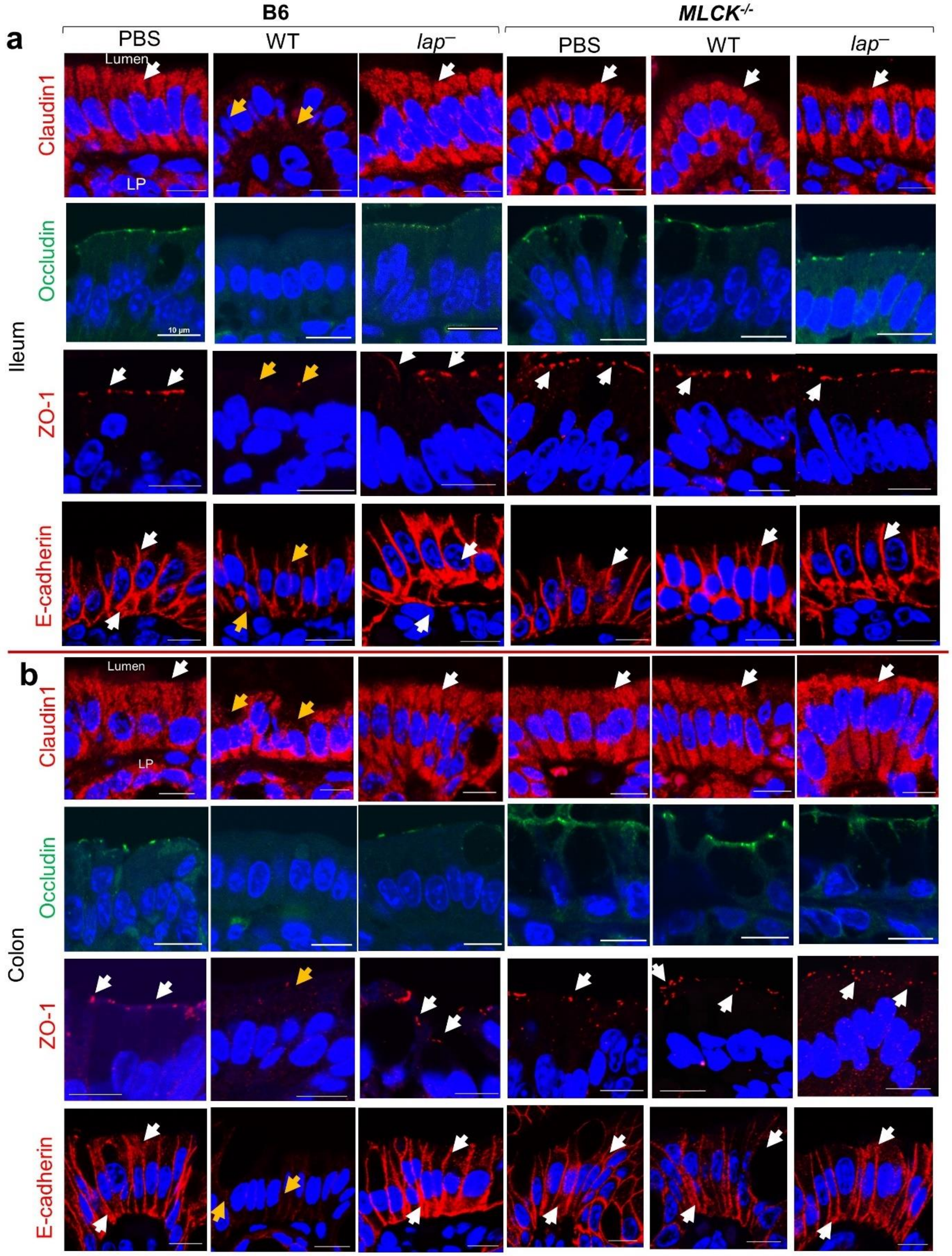


**Supplementary Fig. 15. Tight junction protein profiling.** Representative immunofluorescent images showing the expression and localization of the tight junction proteins Claudin-1(Red), Occludin (Green), ZO-1 (Red), and E-cadherin (Red) in the ileum (a) and colon (b). Nuclei are counterstained with DAPI (blue).

**Supplementary Spreadsheet 1**. Whole genome sequencing comparison of serovar 4b *Listeria monocytogenes* gastroenteritis outbreak strains (H7014 and F4244)
